# GENOMICON-Seq: A comprehensive tool for the simulation of mutations in amplicon and whole exome sequencing

**DOI:** 10.1101/2024.08.14.607907

**Authors:** Milan S. Stosic, Jean-Marc Costanzi, Ole Herman Ambur, Trine B. Rounge

## Abstract

GENOMICON-Seq is a comprehensive genomic sequencing simulation tool that enables the assessment of laboratory and bioinformatics parameters influencing the detection of mutations. The tool generates genomes with mutations, mimicking processes such as low-frequency mutations, APOBEC3 activity in viruses, somatic mutations and single base substitution (SBS) mutational signals. GENOMICON-Seq adds amplicon and whole exome sequencing biases and errors. It outputs sequencing reads compatible with mutation detection tools and a report on mutation origin (generated mutations and PCR errors), nucleotide context, and position. GENOMICON-Seq aids in the evaluation of bioinformatics tools and experimental designs, reducing the need for costly real-world sequencing experiments.

## 1. Background

Advancements in sequencing technologies have revolutionized our ability to detect low-frequency mutations, unveiling a new layer of genetic diversity. In viruses, low-frequency mutations denoted as intrahost single nucleotide variants (iSNVs), can reveal genetic diversity that informs viral evolution and pathogenicity ^1–7^. In humans, the accumulation of subclonal low-frequency somatic mutations is a major cause of cancer development ^8–12^. Somatic mutations vary, and their frequencies are shown to differ in different cancer types ^13^. However, detecting low-frequency mutations presents a significant challenge.

The detection of low-frequency mutations can, however, be accomplished by amplicon sequencing. While powerful, this method introduces potential technical mutations during PCR and sequencing, complicating the mutation detection ^14–16^. Various detection strategies are employed to discriminate between true mutations and technical artefacts, including criteria such as minimum allele frequency, read coverage, and the number of reads supporting a mutation ^6,17–20^. However, these criteria are often subjective derived rather than grounded in empirical data ^17^. Despite the efforts to ensure detection accuracy, standardized and validated methods are lacking ^21^. In this context, simulated datasets could be invaluable in systematically evaluating the reliability of detection strategies.

Whole genome sequencing, along with its less expensive alternative, whole exome sequencing (WES), is employed for detecting low-frequency somatic mutations in humans. In WES, the enrichment of the exonic regions is accomplished by probes, rendering sequencing errors the major source of bias in the detection of somatic mutations ^22,23^. Variant callers such as Mutect2 ^24,25^, Strelka2 ^26^, VarScan ^27^, LoFreq ^28^, and SomaticSniper ^29^ have been developed to address these challenges ^30^. Variant callers are often benchmarked using clinical data or simulated data.

Various simulation tools have been developed, each with distinct features and applications. One approach introduces mutations into the Binary Alignment Map (BAM) files using BAMSurgeon ^31^ or SomatoSim ^32^. A more complex simulator, NEAT ^33^ allows customization of many parameters, aiming to reproduce the properties of real-life sequencing data. However, NEAT does not simulate biological or sequencing processes ^33^. Currently, to our knowledge, no tools explore how biological and sequencing processes affect the detection of low-frequency mutations.

We have developed GENOMICON-Seq (GENOmics Modeling, In-silico CONstruction, and Sequencing, https://github.com/Rounge-lab/GENOMICON-Seq), a simulation tool that models the biological signal of mutations and technical biases and errors introduced during library preparation and sequencing. GENOMICON-Seq supports the simulation of amplicon sequencing and WES with PCR and probe-capturing biases, and sequencing errors. This tool generates sequencing FASTQ data that can be used to evaluate ranges of mutational frequencies reliably differentiated from technical noise. GENOMICON-Seq can, therefore, guide better experimental design and methodological improvements. Here, we introduce GENOMICON-Seq and showcase the tool’s utility by exploring the effects of varying polymerase error rates, sample inputs and sequencing depths on the detection of low-frequency mutations in the amplicon sequencing and WES simulators.

## 2. Results

The GENOMICON-Seq is a comprehensive tool that simulates various biological-signal-to-noise ratios during amplicon sequencing and WES (Figure 1A). In this context, ‘noise’ refers to error mutations introduced during PCR or sequencing simulations. In total, 22 parameters influencing this ratio can be set and evaluated (Figure 1B, see tool manual^34^).

**Figure 1.**
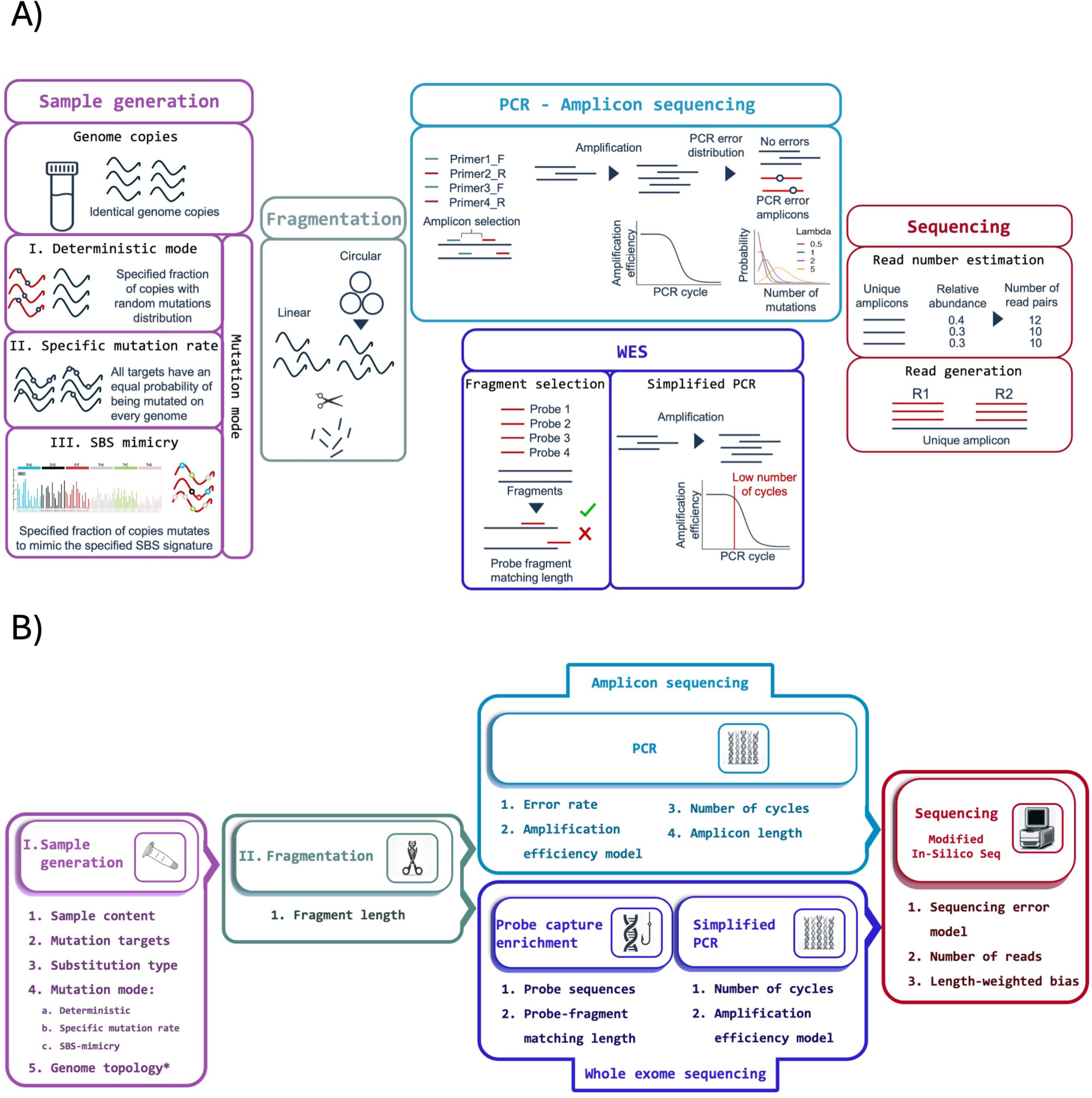
A) Illustration of the GENOMICON-Seq simulated steps. Sample generation includes three mutational modes. Fragmentation provides options for linear and circular genomes. For WES, probe-capture enrichment selects the fragments based on the match between the probe and the fragment, while short, simplified PCR simulation amplifies their numbers following the inverted sigmoid function. In amplicon sequencing, PCR simulation encompasses the mutation introduction after each cycle following the Poisson distributions (illustrated right graph), while the inverted sigmoid function models the amplification efficiency of every cycle (illustrated left graph); the sequencing step employs the error model generator from InSilicoSeq modified to generate reads from each amplified fragment based on their number of copies. B) Overview of GEOMICON-Seq key customizable parameters. Both amplicon and whole exome sequencing (WES) simulations share common processes such as sample generation, fragmentation, and sequencing. PCR enrichment is specific to the amplicon sequencing simulation, while probe-capture enrichment is unique to the WES simulation. *Genome topology, specifying whether the genome is circular or linear, is only relevant in amplicon sequencing and not applicable in whole exome sequencing. Single Base Substitution (SBS)-mimicry mode is applicable only in whole exome sequencing.

Both amplicon and WES simulations start with a mutational signal introduction by “deterministic” (customizable total number of mutations and mutated genomes), “specific mutation rate” (mutations are introduced with a customizable mutation rate), and “SBS (Single Base Substitution)-mimicry” mode (generated mutations followed the specific SBS mutational signal). The amplicon sequencing simulation includes PCR simulation and is adapted from the TaME-seq2^19^ and Cullen et. al ^35^ methods for Human papillomavirus (HPV) whole-genome sequencing. The WES simulation is modelled after the SureSelect whole exome sequencing protocol ^36^ simulating probe-capture enrichment. However, it can be used with any provided probe sequences. Sequencing simulation employs a modified InSilicoSeq tool generating reads with error models from different Illumina instruments ^37^.

GENOMICON-Seq represents a collection of scripts written in R, C++, Python, and Powershell, integrating several well-known tools, orchestrated by Snakemake ^38^, and packed in a Docker container ^39^. It produces FASTQ files and a CSV file with details of the generated mutations. For amplicon sequencing, it also generates a CSV file with information on PCR error mutations.

### 2.1. Study cases

We utilized study cases to demonstrate the application of GENOMICON-Seq. In amplicon sequencing, study cases A1-3 evaluated how generated mutations in TCN context with “specific mutation rate” mode and polymerase error rates (A1), initial genome copy number and the amount of sample input used in the simulated library preparation (A2), and sequencing depth (A3) influence mutation detection. The genome used was HPV 16, a small oncogenic virus with a genome length of 7906 bp ^40^.

In the WES simulation study cases, we utilized the assembled sequence of chromosome 1 (chr1) of the human genome (GRCh38 assembly ^41^). Study cases W1 and W2 illustrated the impact of sequencing depths (20M and 35M generated reads, respectively) on detecting alternative alleles. W3 replicated the parameters of case W2 but incorporated a length-weighted sequencing bias exemplifying its effect on mutation detection. In W1-W3 generated samples, we employed “deterministic” mode generating mutations in TCN context, varying the copy number of chr1 and the number of mutated positions.

Study case W4 utilized the “SBS-mimicry” mode to generate samples with SBS2, SBS8, and SBS24 mutational signatures obtained from the Catalogue of Somatic Mutations in Cancer (COSMIC) ^42^. For each sample, 35M reads were generated without the length-weighted bias.

#### 2.1.1. Amplicon sequencing study cases

The overview of generated reads and mapping statistics are presented in Table S1 for study cases A1-3. In study case A1, as the polymerase error rate increases, a growing proportion of fragments will carry PCR errors. A higher proportion of fragments carrying an error makes the process of assigning read pairs to a fragment more challenging. Consequently, the total number of reads generated deviated from the expected. In addition, a higher number of mutations (generated and polymerase errors) resulted in lower mapping rates indicated by a higher number of unmapped reads (Table S1).

The number of mutated and non-mutated genomes, total number of unique mutations, sample mutation frequency (total number of mutations per Mb DNA in the *in-silico* sample), mean variant allele frequency (VAF, fraction of genome copies harbouring a generated mutation), and the mean number of PCR errors are shown in Table S2. As expected, sample mutation frequency and VAF increased with the increasing mutation rate.

Study case A1 was set up to evaluate how polymerase and sequencing errors influence the detection of generated mutations. At the lowest mutation rate, corresponding to a mutation frequency of 0.6 mutations per Mb DNA, and 0.00001 VAF in the sample (Table S2), the distribution of read counts supporting generated mutations and noise overlapped. The read count supporting generated mutations increased with increasing mutation rates (mutation frequency of ∼ 16 and 488 mutations per Mb DNA and 0.0002-0.006 VAF, Table S2). However, the read counts for both noise and generated mutations positively correlated with the mutation rate (Figure S1A, Table S3). This positive correlation is likely due to mapping errors at high mutation rates.

To exclude all FP (false positives, PCR and/or sequencing errors) detections in non-mutated samples, the necessary read-count cutoff (the number of reads required for calling a variant) ranged between 280 and 485, increasing with increasing polymerase error rate. In mutated samples, we identified the number of TPs (true positives, detected generated mutations), FPs, and FNs (false negatives, generated but not detected mutations) at a range of read-count cutoffs (1-1000, increasing by 5) to determine the optimal read-count cutoff with maximized precision and recall. Mutation rates influenced the optimal read-count cutoff ranging between 10 in lower and over 500 in high mutation scenarios (Figure 2A, Figure S2, Table S4). In addition, at lower mutation rates both precision and recall improved with a higher mutation rate but slightly decreased at the highest polymerase error rate. However, at the highest mutation rate corresponding to 0,006 mean sample-VAF, precision and recall remained stable reaching a maximum of ∼ 0.5 regardless of the polymerase error rate.

**Figure 2.**
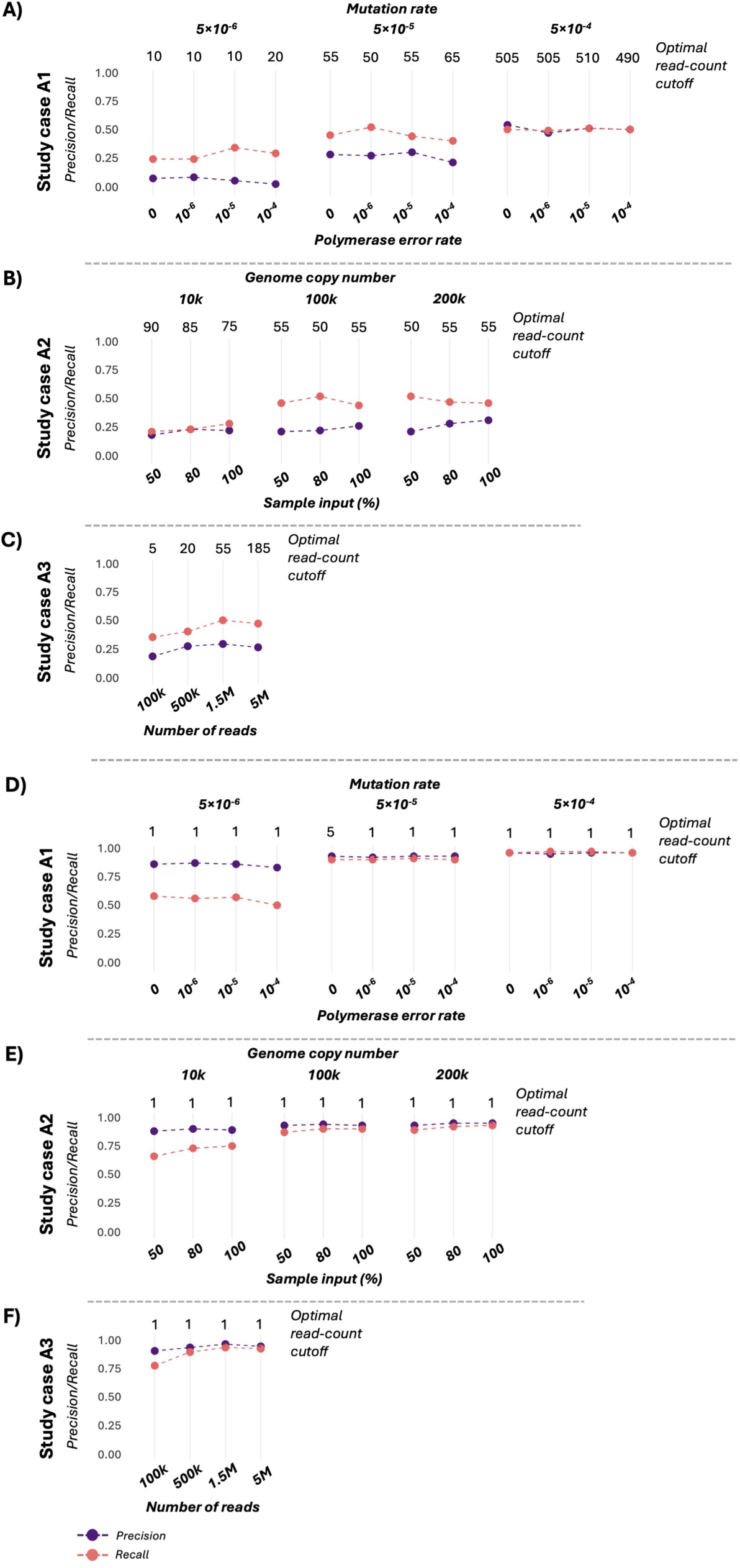
Mean precision (purple) and recall (pink) at the optimal read count cutoff. A) study case A1, B) study case A2 and C) study case A3; and for APOBEC3 signatures in A) study case A1, B) study case A2 and C) study case A3.

In study case A2, we evaluated how the number of initial genome copies and the fraction of sample input used for library preparation influenced the detection of generated mutations. The mutational-signal-to-noise ratio remained almost unchanged due to the constant mutation and polymerase error rate (Figure S1B). The optimal read-count cutoff decreased from 90 to 50, stabilizing between 50-55, with the rising number of genome copies and sample inputs (Figure 2B, Figure S3, Table S5). At low genome copy numbers (10k), precision and recall increased with the increase of sample input. At a high copy number (100k-200k), precision still increased, while recall slightly decreased with the increase of sample input (Figure 2B).

The relationship between sequencing depth and detection of generated mutations was investigated in the study case A3. The signal-to-noise ratio increased with sequencing depth (Figure S1C, Table S3) necessitating a higher optimal read-count cutoff to maximize precision and recall (Figure 2C). At the optimal read-count cutoff, higher sequencing depth generally improved precision and recall. However, 5M generated reads diminished recall compared to samples with 1.5M reads (Figure 2C, Figure S4, Table S6).

To demonstrate how noise masks the detection of generated mutations in specific contexts, we narrowed the analysis to APOBEC3-induced mutation signatures (Figure 2D-F, Figure S5, Table S7). The optimal read-count cutoff was between 1 and 5 in all samples in all study cases while precision and recall were greatly improved.

#### 2.1.2. WES simulation study cases

Capabilities of the GENOMICON-Seq WES simulation were demonstrated in four study cases utilizing the “deterministic” mode of mutation generation. For W1-3 study cases, the mean number of generated reads and the unmapped reads are shown in Table S8, and the percentage of covered exons is presented in Tables S9-11. While 35M reads resulted in almost all exons covered on average with >300x, 20M reads covered approximately all exons between 50x and 100x. The number of mutations, mutation frequency (in terms of the number of mutations per Mb DNA), and mean VAF are presented in Table S12.

Initially, Mutect2 was used for pairwise comparison of technical replicates of non-mutated samples, and no mutations passed the filtering step. In mutated samples across all study cases, the precision of mutation detection ranged between 0.7 and 1 (Figure 3). The lowest precision and recall were identified in samples with the lowest number of mutated positions and genomes. Recall ranged between 0 and 0.9 across all scenarios (Figure 3, Table S13-S15). However, it reached a plateau when the percentage of mutated genomes was between 10 and 20, corresponding to a mutation frequency of ∼ 0.4-0.9 mutations per Mb DNA, and 0.04-0.1 VAF in the sample. However, the high number of generated mutations resulted in a lower recall plateau (Figure 3). The number of TPs in study case W1 (20M reads, Table S13), was lower than in W2 (35M reads, Table S14) which again was lower than in W3 (35M reads and length-weighted bias, Table S15).

**Figure 3.**
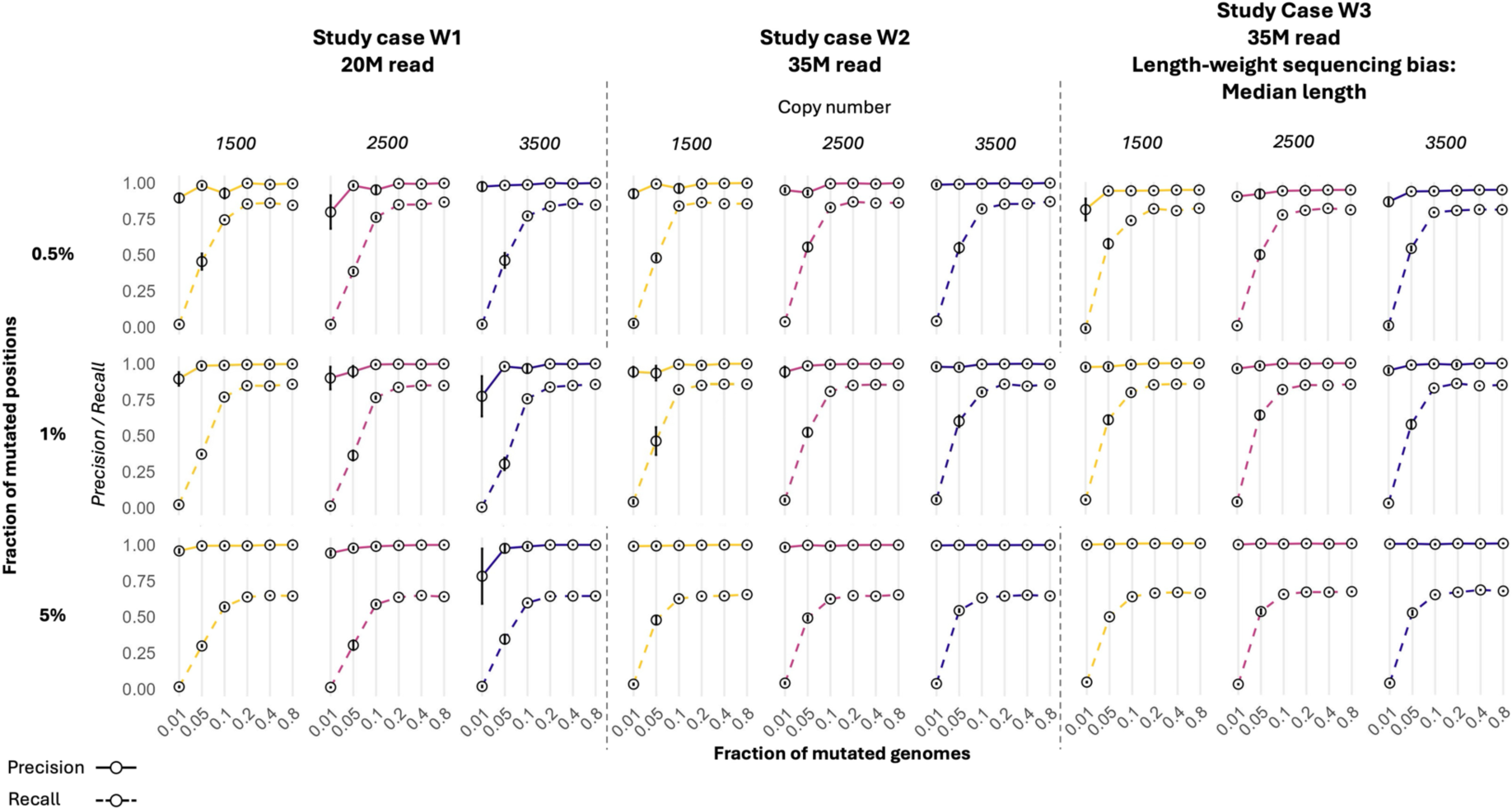
Mean precision and recall across samples in study case W1-W3. Each plot represents a varying fraction of mutated genomes (0.01, 0.05, 0.1, 0.2, 0.4, and 0.8, x-axis) with an initial number of genome copies (1500, 2500, 3500), and a fraction of mutated positions (0.5%, 1%, and 5%). Solid trend lines connect the precision values, while dashed trend lines connect the recall values of each sample. Whisker (vertical lines) indicate the standard error (SE) calculated based on 3 sample replicates.

In study case W4, we employed the extension of the “deterministic” mode, SBS (Single Base Substitution) -mimicry mode. Mapping statistics, the percentage of covered exons and their sequencing depth, and the number of generated mutations are shown in Tables S16-S18, respectively. TPs were categorized into the 96 SBS mutational signatures. The detected signatures were compared to the signature plot from COSMIC ^42^ (Figure 4). More than 50% of all generated mutations were not detected. However, generated SBS signatures, and detected SBS signatures corresponded to the original COSMIC signature plots.

**Figure 4:**
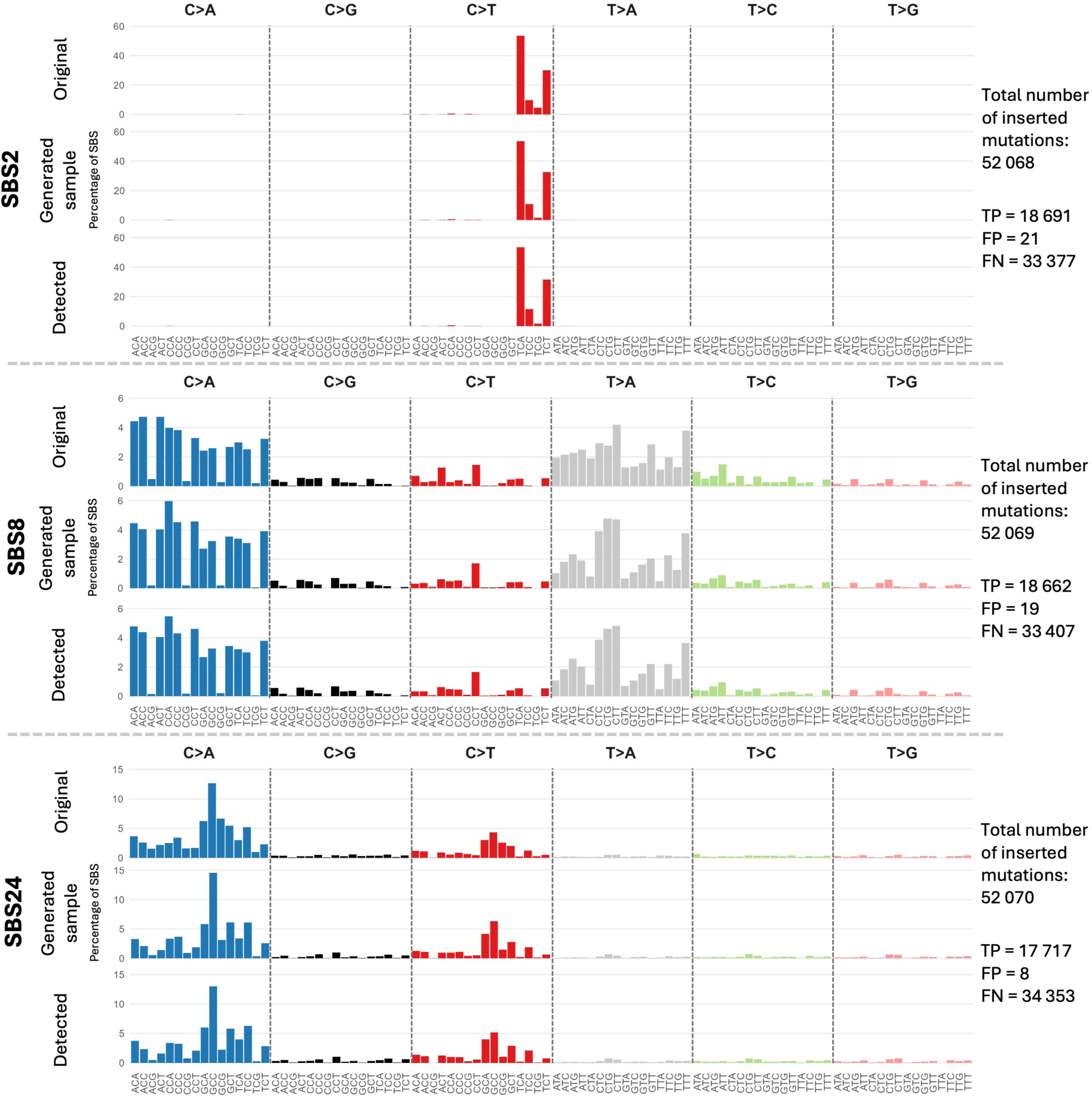
Comparison of generated, detected, and original COSMIC SBS2, 8, 24 mutational signatures. Information about the total number of generated mutations is presented on the side of the plots detailing the number of true positives (TPs), false positives (FPs), and false negatives (FNs).

### 2.2. Performance and processing time

GENOMICON-Seq operates on Unix-like systems requiring only pre-installed Docker but is optimized for high-performance computing (HPC) environments efficiently distributing the workload on multiple cores. Simulations can be run on personal computers provided the workload is scaled down (see tool manual ^34^).

Processing time varies depending on parameters such as polymerase error rate (in amplicon sequencing), mutation mode, genome copies, and genome length. Processing time for one amplicon-sequencing sample replicate with two PCR simulations varied between ∼25 minutes and 4.5 hours per replicate in our study cases with 48 CPUs and 2GB memory per CPU. An increase to 100 CPUs reduced processing time to ∼15-45 minutes per replicate. For the WES simulation involving chr1, the processing time was between ∼55 and 110 minutes per sample with 24 CPUs and 8 GB memory per CPU. The human genome (2500 copies) can be processed with 60 CPUs and 8Gb per CPU in ∼18-24 hours depending on the number of inserted mutations.

## 3. Discussion

GENOMICON-Seq represents a novel approach for genomic sequencing simulation allowing control over mutational processes, library preparation biases, and sequencing errors. This tool supports the simulation of two key library preparation methods, amplicon sequencing and WES. By incorporating the InSilicoSeq ^37^ tool that mimics the sequencing error of Illumina instruments, GENOMICON-Seq produces realistic sequencing reads. GENOMICON-Seq can guide the optimization of sequencing protocols and refine the detection of low-frequency mutations, saving time and resources before committing to expensive real-world sequencing.

Processing time for both amplicon and WES depends on the number of CPUs. PCR simulation in amplicon sequencing is the most resource-demanding process in terms of the number of CPUs. The generated FASTQ files can be processed with any variant calling pipeline, and the mutation information can be used to classify the mutations into TPs, FPs, and FNs.

Amplicon sequencing is commonly employed for analyzing specific genes or small genomes. It can be, for example, used for analyzing whole genome variations in non-culturable viruses. Key determinants of low-frequency mutation detection include the frequency of mutations in the viral sample ^43,44^, polymerase error rate ^16,45^, viral load ^46,47^, sample input in the library preparations ^48,49^, and sequencing depth ^50,51^. Our study cases addressed these factors to demonstrate the GENOMICON-Seq applicability.

HPV16 amplicon sequencing was simulated to replicate the laboratory preparation used in the TaME-seq method ^18,19^. In our study cases, we simulated random APOBEC3 mutation insertion in a viral population by utilizing the “specific mutation rate” mode. In this mode, all targeted positions on all viral copies have an equal likelihood of being mutated. APOBEC3 is known to randomly target specific TCN trinucleotide contexts on viral genomes^52–54^.

The choice of the PCR polymerase can markedly influence error rates ^16,45,55^. We simulated the effect of increasing polymerase error rates in study case A1. Increasing the noise level makes it harder to discriminate between generated mutations and noise. GENOMICON-Seq can be used to test how various polymerase error rates affect low-frequency mutation detection.

In study case A2, a higher number of viral genome copies and sample input in the library preparation improved mutation detection. Sample quality, reflected by the number of genomes, and available sample volume can often be limited in real-world conditions, thereby influencing mutation detection ^48,49^. Using a higher genome copy number in the simulated library preparation enhanced the detection of low-frequency mutations. GENOMICON-Seq could detect the minimum sample input required for optimal mutation detection, minimizing sample usage.

Study case A3 highlighted the critical role of sufficient sequencing depth. Higher sequencing depth has been shown to improve the detection of low-frequency mutations^18,19,50^. However, we showed that simulated very high depth also amplified the noise confounding the mutation detection. GENOMICON-Seq can assist in detecting the optimal sequencing depth.

Furthermore, our amplicon sequencing study cases also illustrated that optimal read count cutoffs for maximizing precision and recall varied substantially under the simulated conditions. However, maximizing precision and recall invariably resulted in the loss of TPs and the inclusion of FPs. Defining specific mutational contexts significantly improved the detection of generated mutations. In HPV16 clinical samples, analyzing specific contexts has revealed a decrease in APOBEC-specific mutations in precancerous cervical lesions compared to normal samples ^56,57^.

The detection of HPV iSNVs indicating intrahost variability has been associated with cervical cancerogenesis ^18,19,35,53,56–59^. Moreover, intrahost variability has also been detected in other viruses such as HIV ^60,61^, SARS-CoV-2 ^62,63^, Influenza ^64,65^, and Hepatitis ^66,67^. Yet, standardized methods for assessing intrahost variability are still lacking ^21^. GENOMICON-Seq can be used to evaluate diverse detection strategies.

In humans, the accumulation of low-frequency mutations, referred to as somatic mutations, has been associated with tumorigenesis ^8–12^. Known factors influencing the detection of somatic mutations in WES are mutation frequency ^68^, choice of probes and their specificity ^69^, sample quality ^70^, sequencing depth ^71,72^, and choice of the variant caller ^25–29,70^. In our WES study cases, we employed the “deterministic” mutation mode, varying the number of mutated genomes and the number of mutated positions. This mode roughly models the heterogeneity of cancer, thus reflecting the complex nature of cancer cell populations.

The study cases exemplified the impact of the sequencing depth and length-weighted bias during sequencing. The sequencing depth is known to impact the detection of low-frequency mutations in WES ^71,72^, also shown in our study cases. Moreover, we introduced a length-weighted bias to replicate the behavior of an Illumina sequencer which tends to favor shorter fragments ^73,74^. The bias primarily influenced the number of TPs but did not significantly impact the overall precision and recall.

We employed GATK Mutect2 ^25^, variant caller, known for its efficient removal of FPs, reaching very high precision even with very low mutation frequency ^25,72^, also shown in our simulations. However, a significant number of the generated mutations remained undetected by Mutect2, as indicated by the lower recall values. These results align with previous evaluations, where a low mutation frequency has been linked to reduced recall ^72^. During probe-enrichment capture simulation, DNA fragments are lost resulting in underrepresentation or complete lost of generated mutations. While we only used Mutect2, other variant callers might yield different results. GENOMICON-Seq can be used in assessing and improving variant callers thereby enhancing the detection of somatic mutations.

In addition to the precise identification of somatic mutations, investigation of mutational signatures in cancer has emerged as an area of interest. SBS signatures represent traces of endogenous and exogenous mutational factors reflecting the underlying mechanisms that drive cancer progression ^13,42,75^. In study case W4, we employed the “SBS-mimicry” mode. Even though more than half of the generated mutations were not detected, likely due to the simulated probe-capture enrichment, the distribution of the detected SBS signatures was preserved. The SBS-mimicry mode can facilitate the evaluation of SBS signature detection. In independent study, GENOMICON-Seq has been used to evaluate StarSignDNA, a mutational signature analyser, which successfully identified GENOMICON-Seq’s SBS signatures from a VCF (Variant Calling Format) file ^76^.

GENOMICON-seq has its limitations. Despite being efficient, its full utilization requires HPC environments. The processing time and memory requirements also depend on the simulated parameters (see user manual ^34^). The mutation models do not account for more complex mutational processes, including rearrangements, insertions and deletions. Modelled library preparation steps provide valuable insights and cost-effective assistance in study design, but they do not capture the full complexity of biological and experimental processes.

## 4. Conclusion

GENOMICON-Seq is an innovative and comprehensive simulation tool for generating genomes with user-definable mutations, errors and biases that mimic the amplicon and WES library preparations and Illumina sequencing. It outputs sequencing reads (FASTQ files) and a detailed description of all mutations including their origin (generated mutations and PCR errors), nucleotide context, and position.

Utilizing GENOMICON-Seq, we showed how various polymerase error rates, sample input requirements, and sequencing depth influenced low-frequency mutation detection. Additionally, GENOMICON-Seq can assist in evaluating detection pipelines and strategies, variant caller tools, and the ability to detect specific mutations and SBS signatures. GENOMICON-Seq provides valuable insights into the detection of low-frequency mutations, enhancing research efficiency and reducing the need for costly real-world sequencing experiments.

## 5. Methods

Amplicon sequencing and WES simulators start with sample generation and fragmentation and conclude with read generation. The amplicon sequencing simulator includes PCR simulation and the WES simulator includes probe-capturing enrichment simulation and short PCR mimicking the indexing PCR (Figure 1).

### 5.1. Sample generation

Sample generation includes the specification of genomes/chromosomes present in the sample and their copy numbers. The user controls mutation targets, substitution types, and mutation modes. Mutation targets can be random or specific nucleotides, trinucleotide, or pentanucleotide contexts. Substitution types can be random, can favor transitions over transversions or vice versa, or can be custom-made.

GENOMICON-Seq has three mutation modes, deterministic, specific mutation rate, and SBS-mimicry mode. The deterministic mode allows specifying the fraction of genome/chromosome copies and positions within them that will be mutated. The specified fraction of the genome/chromosome will receive at least one mutation. The fraction of positions represents the total number of introduced mutations. Positions depend on whether the mutation targets are specified (nucleotides, trinucleotide or pentanucleotide contexts). Mutations will be randomly distributed across the specified number of genomes. This mode can be used to simulate clonal expansion and heterogeneity of cancer.

The specific mutation rate mode uses the simplified Kimuar’s infinite sites model which assumes that mutations are rare events and that the probability of two mutations occurring at the same site is negligible. The mutated positions cannot revert to their previous state ^77^. Accordingly, the Poisson distribution models the mutation occurrence probability in a fixed genome length with a user-customizable mutation rate. Once the number of mutations per genome/chromosome copy is determined, the mutated positions will be sampled from the pool of specified targets. Finally, the substitution type will be assigned based on the user specifications.

SBS-mimicry resembles the deterministic mode in that it necessitates the specification of genome/chromosome fraction to receive a mutation and positions to be mutated. However, this mode requires an SBS signature table from the COSMIC database ^42^ that will be translated into probabilities for each trinucleotide context to mutate and the likelihood of substitution by one of the three possible nucleotides. These probabilities govern mutation generation and distribution across the specified number of genomes/chromosomes.

### 5.2. Genome fragmentation

The fragmentation step allows the user to specify the number of fragmentation replicates and a fraction of the original sample that will be fragmented. In the amplicon sequencing simulation, circular genomes are randomly opened before fragmentation. The rest of the fragmentation process is governed by user-customizable fragment length (see tool manual ^34^).

### 5.3. Library preparation protocols

#### 5.3.1. Amplicon sequencing simulation

Fragmented samples can be split into several simulated PCR reactions with specified primers per reaction. The following assumptions are made for the PCR simulation: i) primer binding is perfect; ii) Amplification efficiency drops during the cycling following the inverted sigmoid function, caused by depletion of reagents, accumulation of by-products, and polymerase degradation; iii) Binomial distribution determines the number of copies produced in each cycle; iv) Polymerase error rate inserts mutations following the simplified infinite site model where all nucleotides can be mutated to any other nucleotide with equal probability; v) Complex artefacts are not generated; and vi) each fragment is independently amplified.

Before cycling, user-specified primers are mapped to reference genomes/chromosomes with Bowtie2 ^78^ (v2.4.4.) and cross-referenced with the fragments. The process identifies all possible amplicons representing a combination of all forward and reverse primers binding to a fragment. Amplicons with a length lower than the user-specified length are discarded.

During each cycle, the inverted sigmoid function models the amplification efficiency, while the Poisson distribution determines the number of error mutations new amplicon copies might get. The calculation across various PCRs is facilitated by three key parameters, the midpoint cycle, k-parameter inverted sigmoid function modelling, and the polymerase error rate (see tool manual ^34^).

The fragment entering PCR is initially split into lead and lag units mimicking the denaturation process. New copies of lead/lag units are produced during each cycle, determined by the binomial distribution with the probability equal to the efficiency of the current cycle. Poisson distribution determines the number of error mutations per newly produced copy of lead/lag units. Lead/lag copies with or without an error mutation have an equal probability of being amplified and getting a new error mutation in each cycle. A user-customizable fraction of all amplicons selected for sequencing is governed by the binomial distribution. The process mimics the sub-sampling of a PCR reaction.

#### 5.3.2. Probe-capture enrichment in WES simulation

Probe sequences in FASTA or BED (Browser Extensible Data) format select fragments with the exonic targets. Selection is based on the customizable minimum overlap between probes and fragments. Both options are based on mapping of fragments and probes to specified genomes/chromosomes with Bowtie2^78^ (v2.4.4.) and determining overlapping lengths with Bedtools^79^ (v2.31.1). A short PCR simulation amplifies the fragments, omitting polymerase errors, and keeping the amplification efficiency model as in amplicon sequencing.

### 5.4. Sequencing

Before sequencing simulation, the relative abundance of all amplicons or WES fragments is determined. The multinomial distribution uses a relative abundance of each fragment as a probability determining the number of read pairs that will be generated from each fragment.

For the read generation, we modified the InSilicoSeq ^37^ (v1.6.0) tool which now generates read pairs from each end of the fragment. The tool is still using the integrated error model generator that inserts the sequencing errors.

An additional feature is the introduction of a length-weighted sequencing bias that skews the probability of a fragment being sequenced. Based on the length of all selected fragments, the probability of being sequenced is determined by an inverted sigmoid function for each fragment length (see tool manual ^34^).

### 5.5. Study-cases and downstream analysis

The genome used in the amplicon sequencing simulation was HPV16 A1 (GenBank id K02718.1, genome sequence was retrieved from PAVE database ^40^). The virus is usually submitted to a PCR enrichment. All amplicon sequencing cases were run on 48 CPUs with 2 GB RAM per CPU.

We have chosen three study cases with different sets of parameters (Table 1). In all study cases, the NovaSeq error model ^37^ integrated into the InSilicoSeq was employed during the read generation. Each sample was processed in five replicates. Mutation targets were TCN trinucleotide contexts with a 3:1 transition vs. transversion rate. Fragment lengths ranged between 250 and 550 bp. As we simulated the TaME-seq method lab preparation step, two separate PCR simulations were run for each replicate with a minimum amplicon length of 200 bp. 0.5% of each PCR reaction was submitted to sequencing simulation corresponding to the approximate amount of PCR product used in the TaME-seq library prep.

**Table 1.**
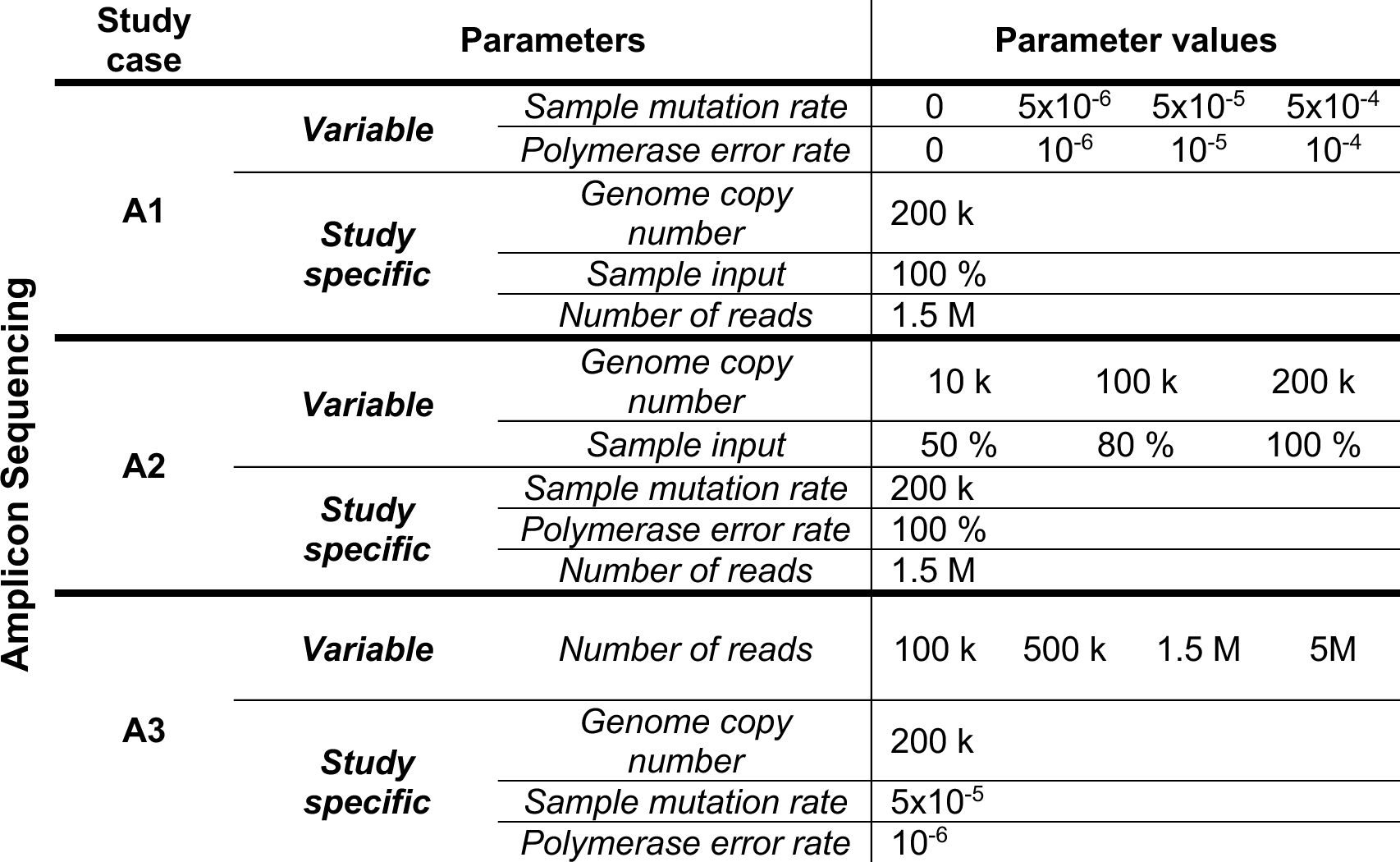
Overview of the parameters used in study cases A1, A2, and A3 of amplicon sequencing simulation.

| Amplicon Sequencing | Study case | Parameters | Parameter values |  |  |  |  |
| --- | --- | --- | --- | --- | --- | --- | --- |
|  | A1 | Variable | Sample mutation rate | 0 | 5x10<sup>-6</sup> | 5x10<sup>-5</sup> | 5x10<sup>-4</sup> |
|  |  |  | Polymerase error rate | 0 | 10<sup>-6</sup> | 10<sup>-5</sup> | 10<sup>-4</sup> |
|  |  | Study specific | Genome copy number | 200 k |  |  |  |
|  |  |  | Sample input | 100 % |  |  |  |
|  |  |  | Number of reads | 1.5 M |  |  |  |
|  | A2 | Variable | Genome copy number | 10 k | 100 k | 200 k |  |
|  |  |  | Sample input | 50 % | 80 % | 100 % |  |
|  |  | Study specific | Sample mutation rate | 200 k |  |  |  |
|  |  |  | Polymerase error rate | 100 % |  |  |  |
|  |  |  | Number of reads | 1.5 M |  |  |  |
|  | A3 | Variable | Number of reads | 100 k | 500 k | 1.5 M | 5M |
|  |  | Study specific | Genome copy number | 200 k |  |  |  |
|  |  |  | Sample mutation rate | 5x10<sup>-5</sup> |  |  |  |
|  |  |  | Polymerase error rate | 10<sup>-6</sup> |  |  |  |

Generated reads from all study cases were processed following the TaME-seq pipeline ^19^ with additional steps outlined in Figure 5A. In short, reads were mapped to the reference (HISAT2^80^, v2.2.1), and the mpileup function from bcftools^81^ (v.1.12) compiled the mapping results at each nucleotide position. The mapping results from two simulated PCR reactions were combined. The alternative allele was called for positions with a mean quality score ≥ 30. We applied a set of fixed read-count cutoffs (1-1000 increasing by 5) to demonstrate their influence on the alternative-allele detection. Due to the known position and substitution type of the introduced mutation, for each fixed cutoff, detected alternative alleles were classified as TPs when matching the introduced mutations or FPs when there was no match. The number of FNs represented the alternative alleles that were introduced but not detected. The precision and recall were calculated for each technical replicate and each fixed read-count cutoff. Finally, the F1 metric was calculated for each cutoff based on the mean precision and recall of each sample enabling identification of the minimum fixed read count necessary to maximize their values.

**Figure 5.**
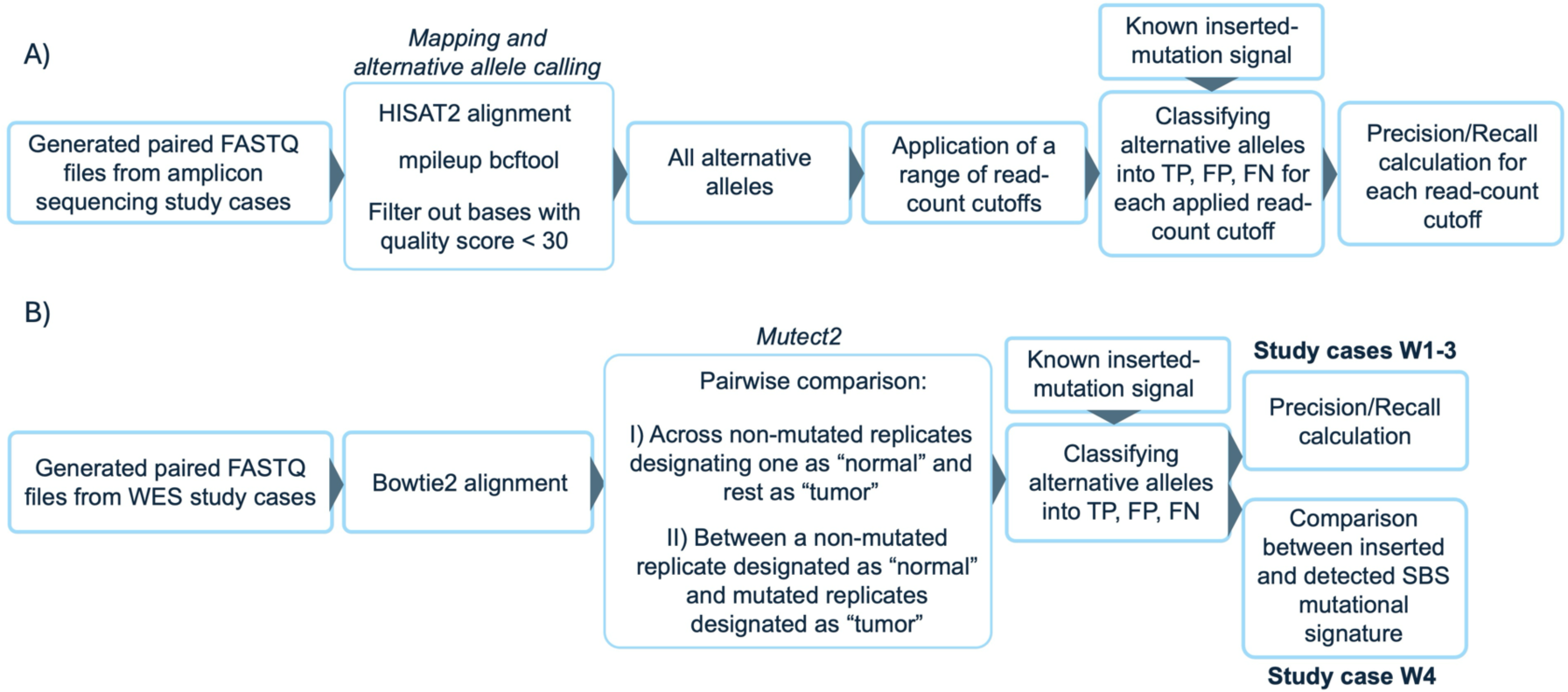
Analysis workflow of the generated FASTQ files in A) amplicon sequencing simulation study cases, and B) WES sequencing simulation study cases.

In the WES simulations, we utilized the assembled sequence of human chr1. Simulations were run on 24 CPUs with 8 GB memory per CPU. The main simulation parameters used in study cases W1-4 are outlined in Table 2. W1-3 simulated samples were processed in three replicates. Generated mutations were in the TCN nucleotide context, with 3:1 transitions vs. transversions ratio. The fragment lengths varied between 250 and 1000 bp. Probe BAM file from xGen™ Exome Research Panel v2 ^82^ was employed in probe-capture enrichment simulation, with a minimum matching length of 100 bp between probe and fragment. The indexing PCR simulation was conducted over eight cycles while the InSIlicoSeq-incorporated NovaSeq error model was used in read generation.

**Table 2.**
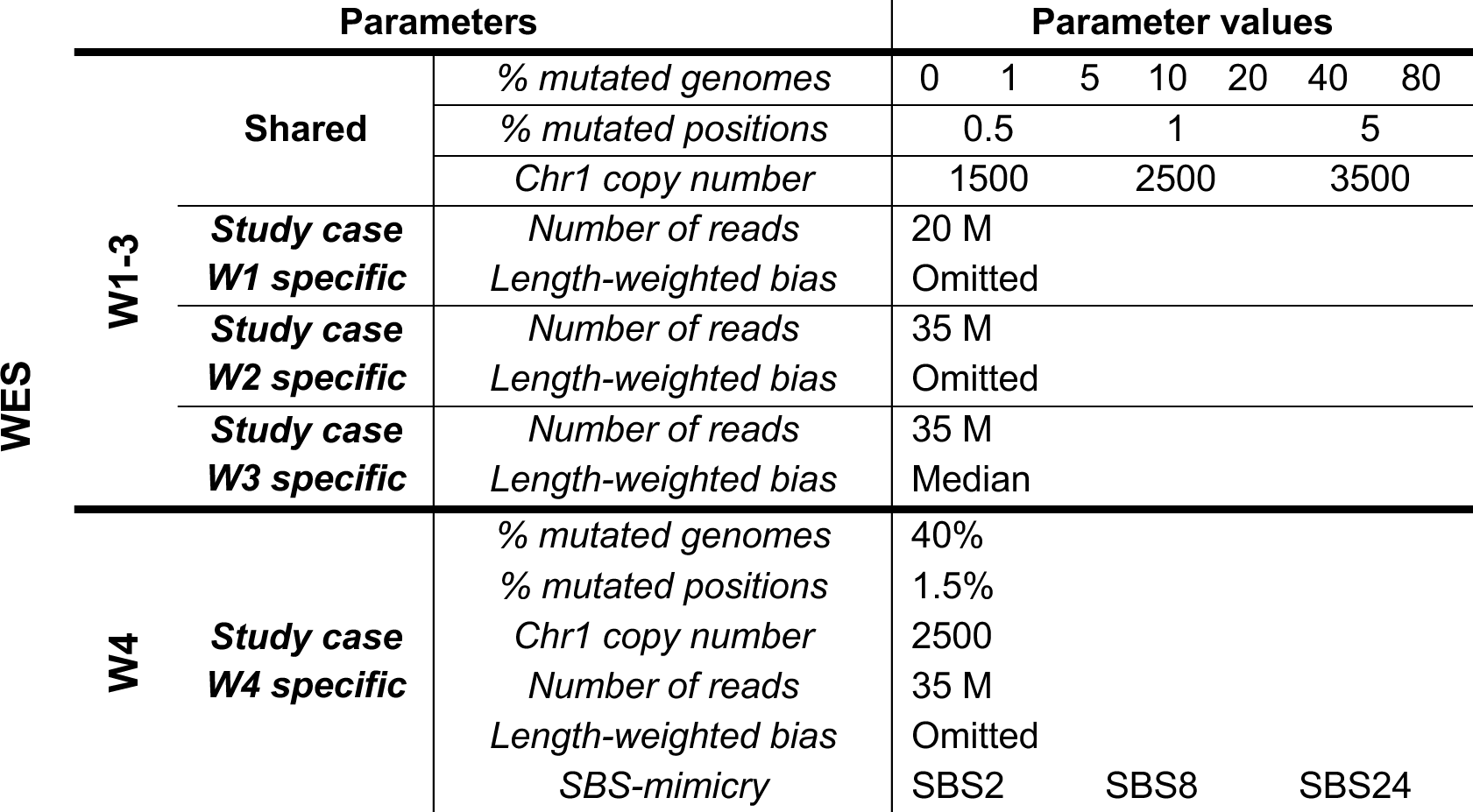
Overview of the parameters used in WES-simulation study cases W1-3, and study case W4.

In the W4 study case, generated samples mimicked SBS2, SBS8, and SBS24 mutational signatures. Out of a total of 2500 chr1 copies, 40% harbored at least one mutation while the total mutation number constituted 1.5% of all positions in the exonic regions (Table 2)

We mapped the generated reads from all WES study cases using Bowtie2 ^78^ (v2.4.4.). Detection of alternative alleles was performed with GATK Mutect2 ^24^ (v4.1.3.0, default settings), as detailed in Figure 5B. We used a replicate of a mutation-free sample as the ‘normal’ control, and mutated samples as ‘tumor’ samples. Additionally, we conducted pairwise comparisons across all mutation-free replicates, designating one as ‘normal’ and the others as ‘tumor.’ Alternative alleles were then filtered with FilterMutectCalls ^83^ (default settings).

Filtered alternative alleles matching the generated mutation’s position and substitution type were classified as TPs, mutations without a match were classified as FPs, while the number of FNs represented the number of generated mutations that were not detected. Finally, for all samples, we calculated the precision and recall in study cases W1-3. In the W4 study case, filtered alternative alleles in all samples were compared to the original and generated SBS signature.

## 6. Ethics approval and consent to participate

Not applicable.

## 7. Consent for publication

Not applicable.

## 8. Availability of data and materials

GENOMICON-Seq is available from https://github.com/Rounge-lab/GENOMICON-Seq/tree/main. The data presented is simulated and can be reproduced by using the specific config files available with the tool.

## 9. Competing interests

The authors declare that they have no competing interests.

## 10. Funding

This work was supported by a PhD grant from the Faculty of Health Sciences, Oslo Metropolitan University. The funders had no role in tool design, in the writing of the report, or in the decision to submit the article for publication.

## 11. Author contribution

TBR and MSS developed the theoretical framework; MSS created the tool; MSS and JMC performed the initial testing and simulation runs; MSS performed variant calling and the analysis of the simulated data; MSS, TBR, and OHA drafted the manuscript text. All authors contributed to the writing and approved the final version of the manuscript.

## 12. Acknowledgements

We would like to thank Irene Kraus Christiansen for her support in establishing the project and valuable insights, and Einar Birkeland for valuable feedback on the manuscript.

## Notes

### Competing Interest Statement

The authors have declared no competing interest.

