## Supplementary information for "GENOMICON-Seq: A comprehensive tool for the simulation of mutations in amplicon and whole exome sequencing"

### Supplementary figures

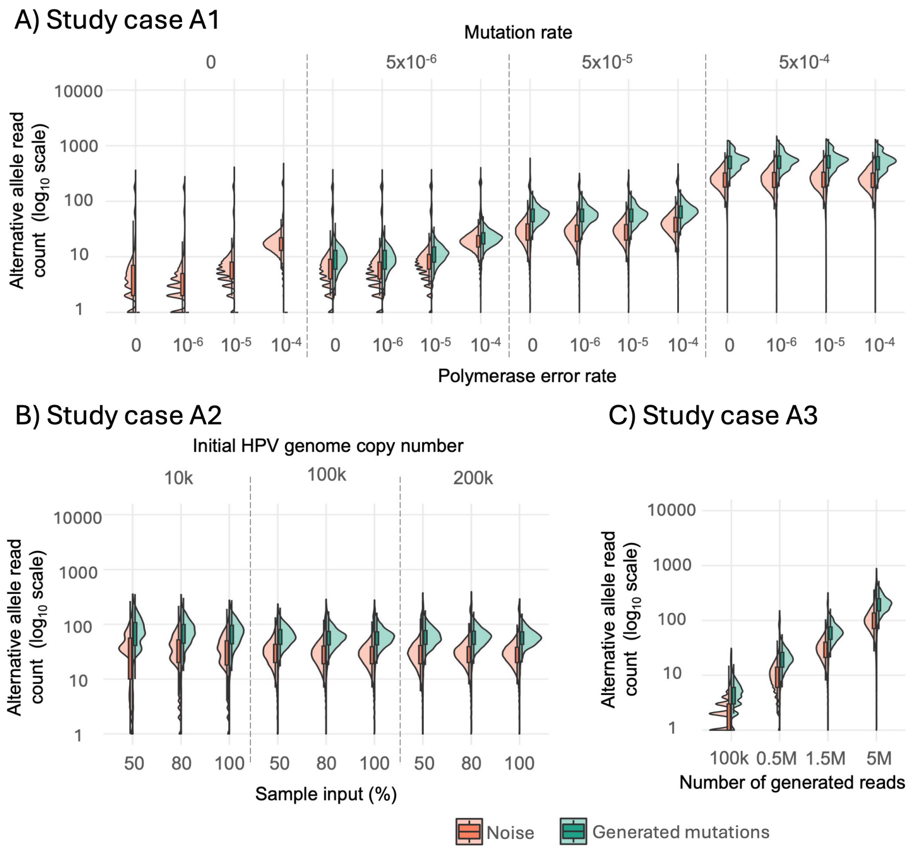


*Figure S1. Distribution of read counts supporting generated mutations and noise in amplicon sequencing simulations in, A) Study case A1, B) Study case A2, C) Study case A3. Split violin plots show the read count distribution of generated mutations and noise on a $log_{10}$-scaled y-axis.*

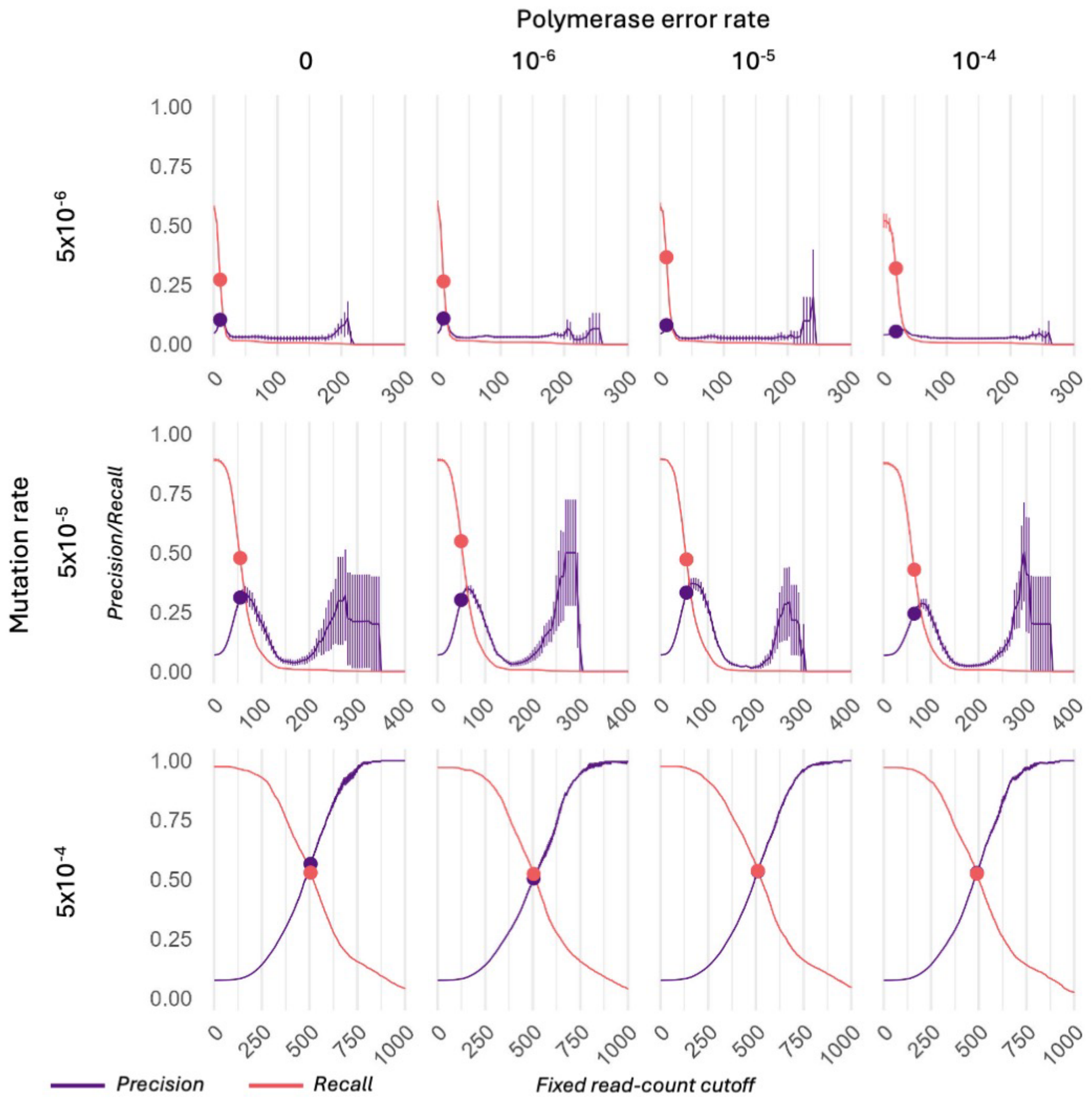


*Figure S2. Mean precision and recall across fixed read-count cutoffs in the study case A1 simulated samples. Each plot in the grid represents a sample generated by varying mutation rates and polymerase error rates. Changes in mean precision (purple) and mean recall (pink), each calculated from five technical replicates of a sample, are plotted against a range of fixed read-count cutoffs on the x-axis. Vertical lines on the graph represent the standard error (SE), calculated from the values across replicates. The x-axis is truncated at the point where both precision and recall drop to zero. Markers on the precision and recall graphs indicate the minimum read-count cutoff where both metrics were maximized and calculated based on the highest F1 value.*

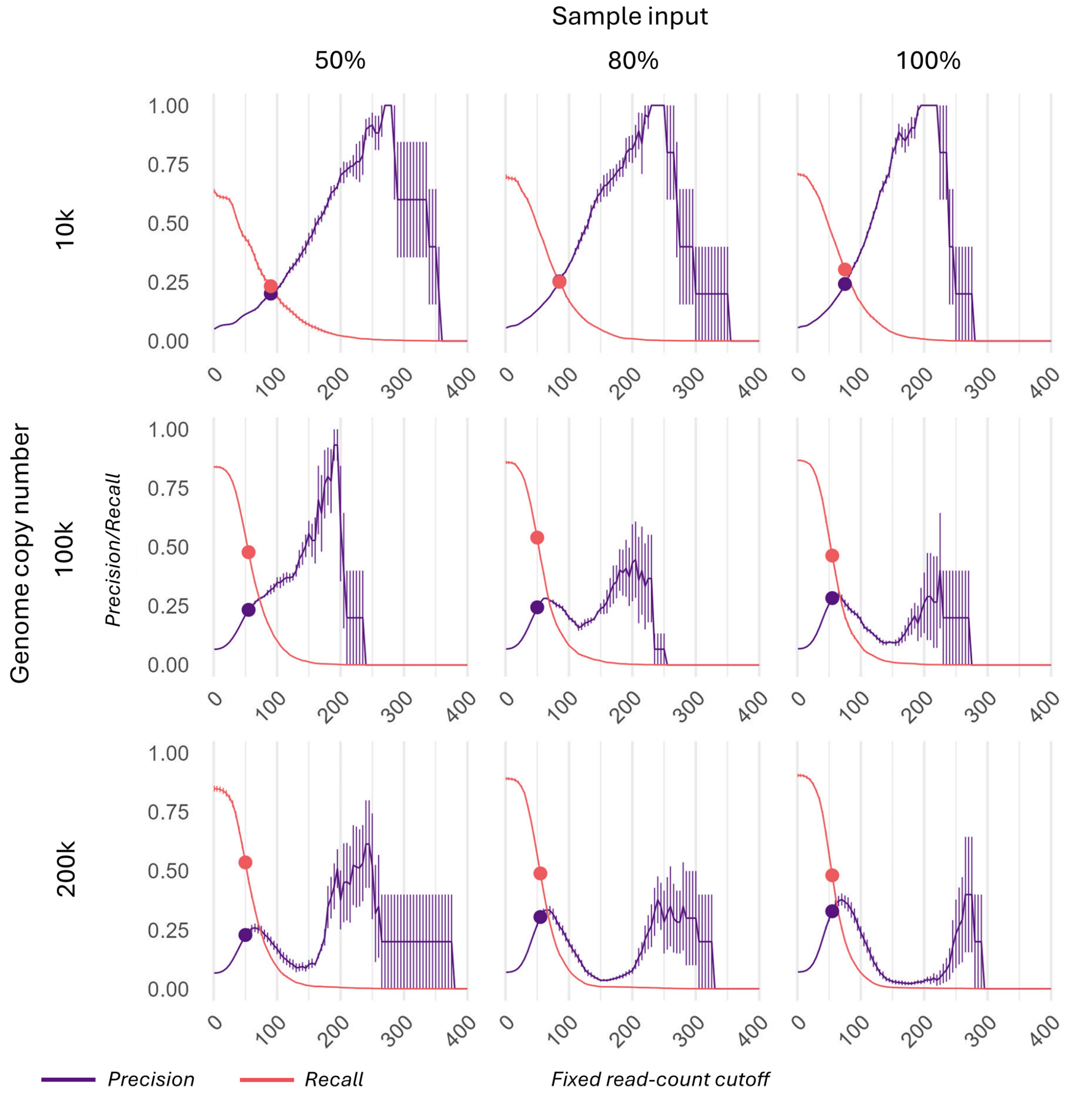


*Figure S3. Mean precision and recall across fixed read-count cutoffs in the study case A2 simulated samples. Each plot in the grid represents a sample generated by varying genome copy numbers and sample input. Changes in mean precision (purple) and mean recall (pink), each calculated from five technical replicates of a sample, are plotted against a range of fixed read-count cutoffs on the x-axis. Vertical lines on the graph represent the standard error (SE), calculated from the values across different replicates. The x-axis is truncated at the point where both precision and recall drop to zero. Markers on the precision and recall graphs indicate the minimum read-count cutoff where both metrics were maximized and calculated based on the highest F1 value.*

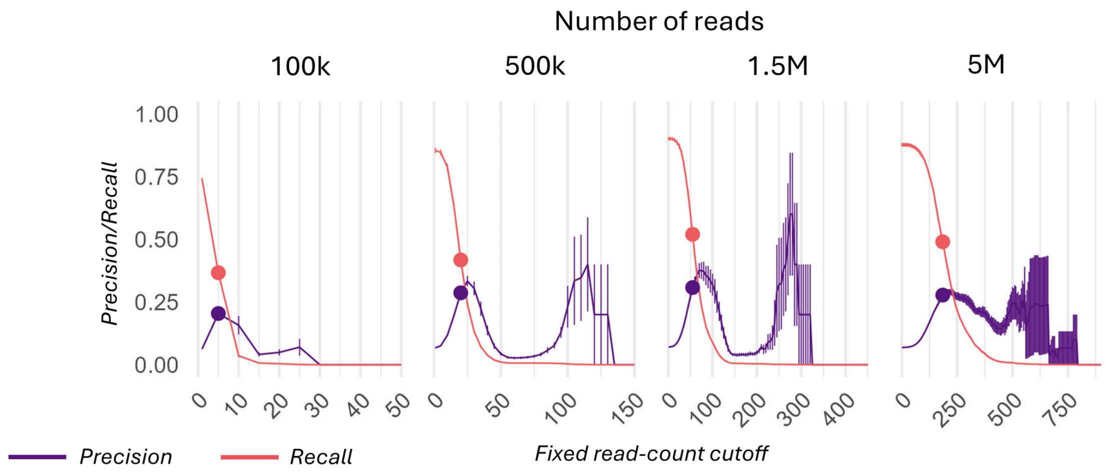


*Figure S4. Mean precision and recall across fixed read-count cutoffs in the study case A3 simulated samples. Each plot in the grid represents a sample generated by varying the number of generated reads. Changes in mean precision (purple) and mean recall (pink), each calculated from five technical replicates of a sample, are plotted against a range of fixed read-count cutoffs on the x-axis. Vertical lines on the graph represent the standard error (SE), calculated from all values across replicates. The x-axis is truncated at the point where both precision and recall drop to zero. Markers on the precision and recall graphs indicate the minimum read-count cutoff where both metrics were maximized and calculated based on the highest F1 value*.

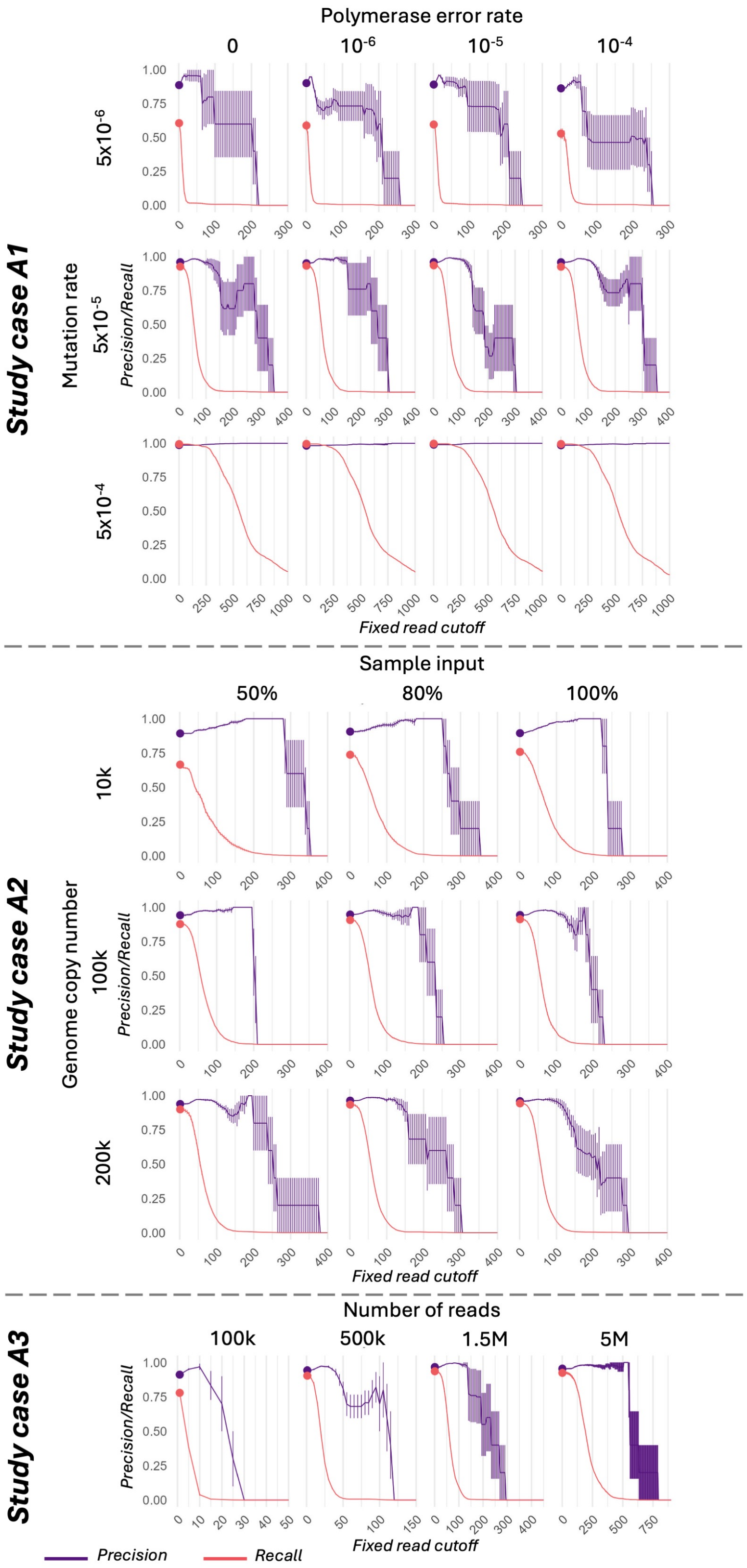


*Figure S5. Mean precision and recall across fixed read-count cutoffs in the study case A1-A3 simulated samples when the analysis is narrowed to APOBEC3 signatures. Each plot in the grid or each study case represents a*

*sample generated by varying mutation rate and polymerase error rate in study case A1, by varying genome copy number and percentage of sample input in study case A2, and by varying the number of generated reads in study case A3. Changes in mean precision (purple) and mean recall (pink), each calculated from five technical replicates of a sample, are plotted against a range of fixed read-count cutoffs on the x-axis. Vertical lines on the graph represent the standard error (SE), calculated from all values across replicates. The x-axis is truncated at the point where both precision and recall drop to zero. Markers on the precision and recall graphs indicate the minimum read-count cutoff where both metrics were maximized and calculated based on the highest F1 value.*

SUPPLEMENTARY TABLES

Table S1.Overview of the mapping statistics from study cases A1-3. The table shows the mean and standard deviation (SD) of the number of generated reads, unmapped reads, coverage and percentage of genome length covered with a minimum of 100x sequencing depth, based on the five replicates of each sample.

**Study case A1**

| Mutation rate | Polymerase error rate | Raw reads | | Unmapped reads | | Coverage | | % genome covered with minimum 100x |
|---|---|---|---|---|---|---|---|---|
| | | Mean | SD | Mean | SD | Mean | SD | |
| 0 | 0 | 1500000.0 | 0.0 | 49.7 | 53.7 | 53530.2 | 69.2 | 100 |
| | $10^{-6}$ | 1499805.2 | 16.4 | 1151.4 | 2013.9 | 53448.9 | 82.4 | 100 |
| | $10^{-5}$ | 1497147.6 | 67.9 | 83.1 | 118.5 | 53465.0 | 57.2 | 100 |
| | $10^{-4}$ | 1406151.6 | 222.6 | 186.5 | 246.6 | 50220.0 | 76.1 | 100 |
| $5\times10^{-6}$ | 0 | 1500000.0 | 0.0 | 557.0 | 1469.5 | 53593.5 | 47.3 | 100 |
| | $10^{-6}$ | 1499816.0 | 16.1 | 47.9 | 35.4 | 53557.7 | 112.0 | 100 |
| | $10^{-5}$ | 1497125.2 | 78.5 | 66.1 | 96.4 | 53419.1 | 65.0 | 100 |
| | $10^{-4}$ | 1406244.0 | 580.0 | 269.2 | 470.3 | 50131.8 | 94.7 | 100 |
| $5\times10^{-5}$ | 0 | 1500000.0 | 0.0 | 97.8 | 111.7 | 53615.8 | 36.1 | 100 |
| | $10^{-6}$ | 1499812.0 | 16.7 | 285.3 | 490.8 | 53557.0 | 87.6 | 100 |
| | $10^{-5}$ | 1497318.4 | 48.9 | 80.6 | 54.9 | 53461.4 | 28.2 | 100 |
| | $10^{-4}$ | 1407024.4 | 443.2 | 158.2 | 150.7 | 50186.6 | 71.6 | 100 |
| $5\times10^{-4}$ | 0 | 1500000.0 | 0.0 | 4570.9 | 814.5 | 53423.1 | 83.7 | 100 |
| | $10^{-6}$ | 1499799.6 | 11.6 | 4669.2 | 1255.0 | 53342.0 | 129.6 | 100 |
| | $10^{-5}$ | 1497205.2 | 72.2 | 5217.5 | 1463.4 | 53217.8 | 90.1 | 100 |
| | $10^{-4}$ | 1406167.6 | 229.9 | 5692.2 | 1024.9 | 49992.3 | 61.9 | 100 |

**Study case A2**

| Genome copy number | Sample input (%) | Raw reads | | Unmapped reads | | Coverage | | % genome covered with minimum 100x |
|---|---|---|---|---|---|---|---|---|
| | | Mean | SD | Mean | SD | Mean | SD | |
| 10k | 50 | 1499840.2 | 21.5 | 185.6 | 104.6 | 53566.4 | 23.2 | 100 |
| | 80 | 1499845.2 | 19.6 | 205.5 | 165.6 | 53558.7 | 14.7 | 100 |
| | 100 | 1499847.4 | 20.3 | 216.4 | 119.8 | 53544.7 | 6.6 | 100 |
| 100k | 50 | 1499827.8 | 23.9 | 173.0 | 223.4 | 53598.2 | 32.1 | 100 |
| | 80 | 1499824.4 | 20.9 | 80.1 | 74.5 | 53566.4 | 65.7 | 100 |
| | 100 | 1499824.4 | 17.4 | 134.2 | 160.6 | 53563.8 | 79.9 | 100 |
| 200k | 50 | 1499823.0 | 17.8 | 132.8 | 128.8 | 53573.0 | 68.5 | 100 |
| | 80 | 1499813.2 | 15.3 | 183.9 | 240.7 | 53590.4 | 110.7 | 100 |
| | 100 | 1499821.4 | 20.8 | 158.4 | 229.0 | 53559.1 | 68.4 | 100 |

**Study case A3**

| Number of reads | Raw reads | | Unmapped reads | | Coverage | | % genome covered with minimum 100x |
|---|---|---|---|---|---|---|---|
| | Mean | SD | Mean | SD | Mean | SD | |
| 100k | 99809.8 | 18.7 | 6.8 | 6.7 | 3570.1 | 8.3 | 100 |
| 500k | 499093.2 | 35.0 | 83.1 | 161.9 | 17809.2 | 48.5 | 100 |
| 1.5M | 1497302.2 | 82.2 | 125.4 | 90.9 | 53530.8 | 64.5 | 100 |
| 5M | 4990953.0 | 125.8 | 371.9 | 171.6 | 178406.1 | 110.8 | 100 |

Table S2. Overview of mutated/non-mutated genomes, and generated and PCR-error mutations. The table shows the number of non-mutated genomes, genomes with at least one inserted mutation (mutated genome), the total number of unique mutations, mutation frequency in terms of the number of mutations per Mb DNA in a sample, variant allele frequency representing the mean number of genome copies carrying the allele divided by the total number of genome copies in the sample, and mean and SD of PCR-error mutations that were sequenced per sample from study cases A1-3. The mean and SD of PCR errors were based on results from two PCR reactions for five replicates for each sample.

| **Study case A1** | | | | | | | | | | |
|---|---|---|---|---|---|---|---|---|---|---|
| **Mutation rate** | **Polymerase error rate** | **Non-mutated genomes** | **Mutated genomes** | | | **Generated mutation** | | | **Sequenced PCR errors** | |
| | | Copy number | Copy number | **Mutations number per genome copy** | | **Total number of unique mutation** | **Mutation frequency** | **Variant allele frequency** | Mean number | SD |
| | | | | Mean | SD | | | | | |
| 0 | 0 | 200000 | 0 | 0.0 | 0.0 | 0 | 0 | 0 | 0.0 | 0.0 |
| | $10^{-6}$ | 200000 | 0 | 0.0 | 0.0 | 0 | 0 | 0 | 4655.6 | 94.7 |
| | $10^{-5}$ | 200000 | 0 | 0.0 | 0.0 | 0 | 0 | 0 | 20195.4 | 33.2 |
| | $10^{-4}$ | 200000 | 0 | 0.0 | 0.0 | 0 | 0 | 0 | 23669.4 | 6.1 |
| $5\times10^{-6}$ | 0 | 192327 | 7673 | 1.2 | 0.4 | 620 | 0.6 | 0,00001 | 0.0 | 0.0 |
| | $10^{-6}$ | 192216 | 7784 | 1.2 | 0.4 | 620 | 0.6 | 0,00001 | 4598.4 | 54.6 |
| | $10^{-5}$ | 192261 | 7739 | 1.2 | 0.4 | 620 | 0.6 | 0,00001 | 20236.0 | 56.7 |
| | $10^{-4}$ | 192480 | 7520 | 1.2 | 0.4 | 620 | 0.6 | 0,00001 | 23678.0 | 9.2 |
| $5\times10^{-5}$ | 0 | 134812 | 65188 | 2.1 | 0.5 | 620 | 16.3 | 0,00021 | 0.0 | 0.0 |
| | $10^{-6}$ | 134666 | 65334 | 2.1 | 0.5 | 620 | 16.4 | 0,00021 | 4613.6 | 34.8 |
| | $10^{-5}$ | 134923 | 65077 | 2.1 | 0.5 | 620 | 16.2 | 0,00021 | 20206.0 | 41.9 |
| | $10^{-4}$ | 134798 | 65202 | 2.1 | 0.4 | 620 | 16.3 | 0,00021 | 23760.8 | 9.9 |
| $5\times10^{-4}$ | 0 | 3730 | 196270 | 4.3 | 1.8 | 620 | 488.8 | 0,00623 | 0.0 | 0.0 |
| | $10^{-6}$ | 3750 | 196250 | 4.3 | 1.8 | 620 | 487.0 | 0,00621 | 4556.6 | 75.6 |
| | $10^{-5}$ | 3777 | 196223 | 4.3 | 1.8 | 620 | 487.9 | 0,00622 | 20280.8 | 59.8 |
| | $10^{-4}$ | 3775 | 196225 | 4.3 | 1.8 | 620 | 488.8 | 0,00623 | 24558.8 | 25.0 |
| **Study case A2** | | | | | | | | | | |
| **Genome copy number** | **Sample input (%)** | **Non-mutated genomes** | **Mutated genomes** | | | **Generated mutation** | | | **Sequenced PCR errors** | |
| | | Copy number | Copy number | **Mutations number per genome copy** | | **Total number of unique mutation** | **Mutation frequency** | **Variant allele frequency** | Mean number | SD |
| | | | | Mean | SD | | | | | |
| 10k | 50 | 6781 | 3219 | 1.6 | 0.6 | 619 | 23.5 | 0,0003 | 4049.2 | 37.9 |
| | 80 | 6705 | 3295 | 1.6 | 0.6 | 619 | 24.3 | 0,0003 | 4215.4 | 71.7 |
| | 100 | 6766 | 3234 | 1.6 | 0.6 | 619 | 23.5 | 0,0003 | 4230.4 | 57.8 |
| 100k | 50 | 67266 | 32734 | 2.0 | 0.5 | 620 | 16.9 | 0,0002 | 4500.2 | 71.0 |
| | 80 | 67233 | 32767 | 2.0 | 0.5 | 620 | 16.9 | 0,0002 | 4542.0 | 70.1 |
| | 100 | 67077 | 32923 | 2.0 | 0.5 | 620 | 17.1 | 0,0002 | 4588.4 | 72.5 |
| 200k | 50 | 134690 | 65310 | 2.1 | 0.5 | 620 | 16.5 | 0,0002 | 4566.6 | 84.1 |
| | 80 | 134486 | 65514 | 2.1 | 0.5 | 620 | 16.5 | 0,0002 | 4623.2 | 56.9 |
| | 100 | 134814 | 65186 | 2.1 | 0.5 | 620 | 16.3 | 0,0002 | 4582.4 | 67.9 |
| **Study case A3** | | | | | | | | | | |
| **Number of reads** | | **Non-mutated genomes** | **Mutated genomes** | | | **Generated mutation** | | | **Sequenced PCR errors** | |
| | | Copy number | Copy number | **Mutations number per genome copy** | | **Total number of unique mutation** | **Mutation frequency** | **Variant allele frequency** | Mean number | SD |
| | | | | Mean | SD | | | | | |
| 100k | | 134988 | 65012 | 2.1 | 0.5 | 620 | 16.4 | 0,0002 | 3116.4 | 60.3 |
| 500k | | 134770 | 65230 | 2.1 | 0.5 | 620 | 16.1 | 0,0002 | 11820.2 | 96.2 |
| 1.5M | | 134702 | 65298 | 2.1 | 0.5 | 620 | 16.4 | 0,0002 | 20235.4 | 54.7 |
| 5M | | 134739 | 65261 | 2.1 | 0.4 | 620 | 16.4 | 0,0002 | 23516.8 | 15.2 |

Table S3. The mean number of alternative alleles per sample with a minimum one read supporting them classified as either noise or inserted mutation, and read count distribution (mean, standard deviation (SD), minimum and maximum) of alternative alleles classified as either noise or inserted mutations based on the results from five technical replicates for each sample in study cases A1-3.

**Study case A1**

| Sample index | Mutation rate | Polymerase error rate | Noise Count | Noise Read count supporting alternative allele Mean | SD | Min | Max | Generated mutations Count | Generated mutations Read count supporting alternative allele Mean | SD | Min | Max |
|---|---|---|---|---|---|---|---|---|---|---|---|---|
| 1 | 0 | 0 | 6706.6 | 11.3 | 30.5 | 1 | 364 | 0 | 0 | 0 | 0 | 0 |
| 2 | 0 | $10^{-6}$ | 6710.4 | 9.0 | 27.1 | 1 | 279 | 0 | 0 | 0 | 0 | 0 |
| 3 | 0 | $10^{-5}$ | 7787.8 | 11.0 | 27.2 | 1 | 414 | 0 | 0 | 0 | 0 | 0 |
| 4 | 0 | $10^{-4}$ | 7894.0 | 22.0 | 29.3 | 1 | 482 | 0 | 0 | 0 | 0 | 0 |
| 5 | $5\times10^{-6}$ | 0 | 7198.4 | 11.0 | 24.9 | 1 | 372 | 357.2 | 12.7 | 20.6 | 1 | 216 |
| 6 | $5\times10^{-6}$ | $10^{-6}$ | 7248.6 | 11.5 | 28.2 | 1 | 369 | 363.2 | 13.0 | 23.5 | 1 | 259 |
| 7 | $5\times10^{-6}$ | $10^{-5}$ | 7465.2 | 13.5 | 27.6 | 1 | 372 | 359.2 | 14.5 | 20.7 | 1 | 241 |
| 8 | $5\times10^{-6}$ | $10^{-4}$ | 7573.0 | 24.0 | 28.9 | 1 | 405 | 323.2 | 25.4 | 24.2 | 2 | 261 |
| 9 | $5\times10^{-5}$ | 0 | 7331.4 | 34.3 | 31.6 | 1 | 597 | 552 | 61.6 | 31.2 | 6 | 345 |
| 10 | $5\times10^{-5}$ | $10^{-6}$ | 7334.8 | 32.5 | 27.9 | 1 | 319 | 552 | 60.4 | 29.3 | 7 | 301 |
| 11 | $5\times10^{-5}$ | $10^{-5}$ | 7334.0 | 33.1 | 26.5 | 1 | 368 | 553.8 | 60.5 | 27.6 | 3 | 302 |
| 12 | $5\times10^{-5}$ | $10^{-4}$ | 7353.4 | 45.2 | 36.6 | 1 | 473 | 543.2 | 69.2 | 31.6 | 9 | 352 |
| 13 | $5\times10^{-4}$ | 0 | 7300.0 | 259.8 | 113.7 | 1 | 871 | 604.6 | 546.1 | 215.0 | 78 | 1243 |
| 14 | $5\times10^{-4}$ | $10^{-6}$ | 7300.8 | 263.2 | 121.0 | 1 | 998 | 602.6 | 547.7 | 214.3 | 81 | 1492 |
| 15 | $5\times10^{-4}$ | $10^{-5}$ | 7296.4 | 265.8 | 119.5 | 1 | 952 | 605.2 | 554.4 | 222.0 | 90 | 1324 |
| 16 | $5\times10^{-4}$ | $10^{-4}$ | 7301.8 | 257.5 | 113.8 | 1 | 920 | 602.8 | 528.6 | 208.8 | 70 | 1293 |

**Study case A2**

| Sample index | Genome copy number | Sample input (%) | Noise Count | Noise Read count supporting alternative allele Mean | SD | Min | Max | Inserted mutations Count | Inserted mutations Read count supporting alternative allele Mean | SD | Min | Max |
|---|---|---|---|---|---|---|---|---|---|---|---|---|
| 1 | 10k | 50 | 7321.0 | 39.1 | 32.6 | 1 | 267 | 393.8 | 82.0 | 53.3 | 1 | 358 |
| 2 | 10k | 80 | 7340.8 | 37.7 | 27.1 | 1 | 226 | 431.4 | 76.6 | 43.3 | 1 | 351 |
| 3 | 10k | 100 | 7340.0 | 36.1 | 24.9 | 1 | 192 | 438.4 | 74.0 | 41.0 | 1 | 277 |
| 4 | 100k | 50 | 7352.0 | 34.1 | 21.3 | 1 | 203 | 521.0 | 64.3 | 30.0 | 1 | 236 |
| 5 | 100k | 80 | 7337.6 | 32.8 | 23.5 | 1 | 288 | 532.8 | 61.8 | 30.1 | 3 | 250 |
| 6 | 100k | 100 | 7343.2 | 33.1 | 24.9 | 1 | 279 | 538.2 | 61.3 | 28.7 | 4 | 273 |
| 7 | 200k | 50 | 7355.2 | 34.8 | 27.5 | 1 | 279 | 526.8 | 63.3 | 31.7 | 1 | 376 |
| 8 | 200k | 80 | 7326.2 | 34.5 | 28.3 | 1 | 395 | 553.0 | 62.1 | 30.0 | 3 | 326 |
| 9 | 200k | 100 | 7320.6 | 32.9 | 26.3 | 1 | 264 | 561.6 | 60.4 | 27.0 | 5 | 292 |

**Study case A3**

| Sample index | Number of reads | Noise Count | Noise Read count supporting alternative allele Mean | SD | Min | Max | Inserted mutations Count | Inserted mutations Read count supporting alternative allele Mean | SD | Min | Max |
|---|---|---|---|---|---|---|---|---|---|---|---|
| 1 | 100k | 6793.6 | 2.9 | 2.7 | 1 | 31 | 458.0 | 4.9 | 2.8 | 1 | 29 |
| 2 | 500k | 7335.4 | 11.9 | 12.2 | 1 | 151 | 529.6 | 21.1 | 11.4 | 1 | 134 |
| 3 | 1.5M | 7329.6 | 34.2 | 25.9 | 1 | 280 | 558.4 | 62.9 | 27.9 | 8 | 320 |
| 4 | 5M | 7350.6 | 114.5 | 72.1 | 1 | 885 | 543.6 | 208.2 | 87.5 | 5 | 791 |

Table S4. Overview of mean precision and recall, mean and SD of true positives (TP), false positives (FP), and false negatives (FN) in all mutated samples (mutation rate ≠ 0) at the most optimal read-count cutoff in study case A1.

| Mutation rate | Polymerase error rate | Optimal read cutoff | Mean Precision | Mean Recall | TP | | FP | | FN | |
|---|---|---|---|---|---|---|---|---|---|---|
| | | | | | Mean | SD | Mean | SD | Mean | SD |
| $5\times10^{-6}$ | 0 | 10 | 0.10 | 0.27 | 169.40 | 33.21 | 1526.60 | 562.04 | 450.60 | 33.21 |
| | $10^{-6}$ | 10 | 0.11 | 0.27 | 165.00 | 25.30 | 1341.60 | 173.01 | 455.00 | 25.30 |
| | $10^{-5}$ | 10 | 0.08 | 0.37 | 227.80 | 17.05 | 2620.00 | 491.58 | 392.20 | 17.05 |
| | $10^{-4}$ | 20 | 0.05 | 0.32 | 199.00 | 29.15 | 3480.80 | 368.06 | 421.00 | 29.15 |
| $5\times10^{-5}$ | 0 | 55 | 0.31 | 0.48 | 296.20 | 12.17 | 667.00 | 125.79 | 323.80 | 12.17 |
| | $10^{-6}$ | 50 | 0.30 | 0.55 | 340.00 | 13.96 | 789.60 | 54.47 | 280.00 | 13.96 |
| | $10^{-5}$ | 55 | 0.33 | 0.47 | 292.40 | 12.62 | 595.80 | 106.06 | 327.60 | 12.62 |
| | $10^{-4}$ | 65 | 0.24 | 0.43 | 265.80 | 15.27 | 826.80 | 52.13 | 354.20 | 15.27 |
| $5\times10^{-4}$ | 0 | 505 | 0.57 | 0.53 | 328.20 | 7.56 | 251.80 | 16.65 | 291.80 | 7.56 |
| | $10^{-6}$ | 505 | 0.50 | 0.52 | 324.80 | 8.41 | 319.20 | 23.82 | 295.20 | 8.41 |
| | $10^{-5}$ | 510 | 0.54 | 0.54 | 333.20 | 5.26 | 289.40 | 11.33 | 286.80 | 5.26 |
| | $10^{-4}$ | 490 | 0.53 | 0.53 | 325.80 | 3.70 | 291.20 | 18.31 | 294.20 | 3.70 |

Table S5. Overview of mean precision and recall, mean and SD of true positives (TP), false positives (FP), and false negatives (FN) in all mutated samples (mutation rate ≠ 0) at the most optimal read cutoff in study case A2.

| Genome copy number | Sample input (%) | Optimal read cutoff | Mean Precision | Mean Recall | TP | | FP | | FN | |
|---|---|---|---|---|---|---|---|---|---|---|
| | | | | | Mean | SD | Mean | SD | Mean | SD |
| 10k | 50 | 90 | 0.20 | 0.23 | 144.00 | 15.33 | 571.00 | 34.86 | 475.00 | 15.33 |
| | 80 | 85 | 0.25 | 0.25 | 155.40 | 7.83 | 458.60 | 26.50 | 463.60 | 7.83 |
| | 100 | 75 | 0.24 | 0.30 | 187.80 | 10.31 | 588.80 | 28.26 | 431.20 | 10.31 |
| 100k | 50 | 55 | 0.23 | 0.48 | 296.60 | 5.98 | 973.00 | 22.48 | 323.40 | 5.98 |
| | 80 | 50 | 0.24 | 0.54 | 335.00 | 7.21 | 1036.80 | 44.06 | 285.00 | 7.21 |
| | 100 | 55 | 0.28 | 0.46 | 287.60 | 16.13 | 732.20 | 95.35 | 332.40 | 16.13 |
| 200k | 50 | 50 | 0.23 | 0.54 | 332.60 | 19.88 | 1125.20 | 54.93 | 287.40 | 19.88 |
| | 80 | 55 | 0.30 | 0.49 | 303.60 | 7.47 | 694.80 | 55.06 | 316.40 | 7.47 |
| | 100 | 55 | 0.33 | 0.48 | 298.60 | 12.48 | 612.80 | 69.75 | 321.40 | 12.48 |

Table S6. Overview of mean precision and recall, mean and SD of true positives (TP), false positives (FP), and false negatives (FN) in all mutated samples (mutation rate ≠ 0) at the most optimal read cutoff in study case A3.

| Number of reads | Optimal read cutoff | Mean Precision | Mean Recall | TP | | FP | | FN | |
|---|---|---|---|---|---|---|---|---|---|
| | | | | Mean | SD | Mean | SD | Mean | SD |
| 100k | 5 | 0.20 | 0.37 | 227.40 | 9.69 | 889.40 | 59.43 | 392.60 | 9.69 |
| 500k | 20 | 0.29 | 0.42 | 259.00 | 13.36 | 648.20 | 73.17 | 361.00 | 13.36 |
| 1.5M | 55 | 0.31 | 0.52 | 322.20 | 11.08 | 729.80 | 83.09 | 297.80 | 11.08 |
| 5M | 185 | 0.28 | 0.49 | 303.80 | 12.07 | 794.40 | 88.19 | 316.20 | 12.07 |

Table S7. Overview of mean precision and recall, mean and SD of true positives (TP), false positives (FP), and false negatives (FN) in all mutated samples (mutation rate ≠ 0) at the most optimal read cutoff in study cases A1-3 when their calculation was narrowed to APOBEC3 induced signal.

**Study case A1**

| Mutation rate | Polymerase error rate | Optimal read cutoff | Mean Precision | Mean Recall | TP | | FP | | FN | |
|---|---|---|---|---|---|---|---|---|---|---|
| | | | | | Mean | SD | Mean | SD | Mean | SD |
| $5\times10^{-6}$ | 0 | 1 | 0.89 | 0.61 | 288.20 | 11.95 | 37.00 | 11.47 | 186.80 | 11.95 |
| | $10^{-6}$ | 1 | 0.90 | 0.59 | 274.60 | 38.28 | 31.20 | 14.20 | 191.40 | 38.28 |
| | $10^{-5}$ | 1 | 0.89 | 0.60 | 282.20 | 28.77 | 34.60 | 7.09 | 190.80 | 28.77 |
| | $10^{-4}$ | 1 | 0.86 | 0.53 | 245.60 | 40.72 | 39.40 | 10.92 | 218.40 | 40.72 |
| $5\times10^{-5}$ | 0 | 5 | 0.96 | 0.93 | 421.40 | 10.01 | 18.00 | 1.73 | 33.60 | 10.01 |
| | $10^{-6}$ | 1 | 0.95 | 0.93 | 416.80 | 7.73 | 21.60 | 5.03 | 30.20 | 7.73 |
| | $10^{-5}$ | 1 | 0.96 | 0.94 | 422.00 | 6.12 | 18.00 | 3.46 | 29.00 | 6.12 |
| | $10^{-4}$ | 1 | 0.96 | 0.93 | 428.40 | 6.02 | 18.60 | 4.39 | 34.60 | 6.02 |
| $5\times10^{-4}$ | 0 | 1 | 0.99 | 0.99 | 459.40 | 0.89 | 6.00 | 2.00 | 2.60 | 0.89 |
| | $10^{-6}$ | 1 | 0.98 | 1.00 | 459.00 | 1.00 | 8.00 | 0.00 | 2.00 | 1.00 |
| | $10^{-5}$ | 1 | 0.99 | 1.00 | 467.80 | 0.45 | 4.60 | 2.30 | 1.20 | 0.45 |
| | $10^{-4}$ | 1 | 0.99 | 0.99 | 473.60 | 1.14 | 6.80 | 1.64 | 2.40 | 1.14 |

**Study case A2**

| Genome copy number | Sample input (%) | Best read cutoff | Mean Precision | Mean Recall | TP | | FP | | FN | |
|---|---|---|---|---|---|---|---|---|---|---|
| | | | | | Mean | SD | Mean | Mean | SD | Mean |
| 10k | 50 | 1 | 0.89 | 0.67 | 318.60 | 8.29 | 38.20 | 1.30 | 160.40 | 8.29 |
| | 80 | 1 | 0.91 | 0.74 | 351.00 | 12.94 | 36.40 | 5.32 | 125.00 | 12.94 |
| | 100 | 1 | 0.90 | 0.76 | 356.40 | 7.02 | 41.80 | 3.11 | 113.60 | 7.02 |
| 100k | 50 | 1 | 0.94 | 0.88 | 404.60 | 4.93 | 25.00 | 7.14 | 56.40 | 4.93 |
| | 80 | 1 | 0.95 | 0.91 | 420.80 | 6.57 | 23.40 | 3.21 | 43.20 | 6.57 |
| | 100 | 1 | 0.94 | 0.91 | 423.60 | 5.03 | 25.20 | 2.39 | 40.40 | 5.03 |
| 200k | 50 | 1 | 0.94 | 0.90 | 425.80 | 12.91 | 27.40 | 4.83 | 47.20 | 12.91 |
| | 80 | 1 | 0.96 | 0.93 | 445.00 | 9.08 | 16.60 | 3.05 | 31.00 | 9.08 |
| | 100 | 1 | 0.96 | 0.94 | 447.40 | 4.16 | 18.60 | 6.43 | 26.60 | 4.16 |

**Study case A3**

| Number of reads | Best read cutoff | Mean Precision | Mean Recall | TP | | FP | | FN | |
|---|---|---|---|---|---|---|---|---|---|
| | | | | Mean | SD | Mean | Mean | SD | Mean |
| 100k | 1 | 0.91 | 0.78 | 371.00 | 7.04 | 35.80 | 3.11 | 105.00 | 7.04 |
| 500k | 1 | 0.94 | 0.90 | 414.20 | 9.98 | 25.00 | 4.30 | 43.80 | 9.98 |
| 1.5M | 1 | 0.97 | 0.94 | 446.80 | 7.09 | 15.60 | 4.98 | 30.20 | 7.09 |
| 5M | 1 | 0.95 | 0.93 | 430.20 | 8.93 | 20.40 | 4.72 | 34.80 | 8.93 |

Table S8. Number of generated and unmapped reads per sample in WES study cases W1-3.

| | Chr1 copy number | Fraction of mutated copies | Raw reads | | Percentage of mutated positions | | | | | |
|---|---|---|---|---|---|---|---|---|---|---|
| | | | | | 0.5% | | 1% | | 5% | |
| | | | | | Unmapped reads | | | | | |
| | | | Mean | SD | Mean | SD | Mean | SD | Mean | SD |
| **Study case W1** 20M reads Length-weighted bias omitted | 1500 | 0.01 | 20000000 | 0 | 3829.7 | 1226.7 | 3523.3 | 1566.8 | 2341.0 | 479.7 |
| | | 0.05 | 20000000 | 0 | 3171.0 | 518.5 | 2430.7 | 950.2 | 2882.7 | 1474.9 |
| | | 0.1 | 20000000 | 0 | 2113.0 | 142.5 | 3453.3 | 372.8 | 2479.7 | 1496.8 |
| | | 0.2 | 20000000 | 0 | 2223.7 | 259.8 | 3574.0 | 2111.5 | 3311.7 | 506.1 |
| | | 0.4 | 20000000 | 0 | 1901.3 | 343.6 | 2638.0 | 558.5 | 2924.3 | 1567.7 |
| | | 0.8 | 20000000 | 0 | 2869.0 | 829.1 | 1918.7 | 494.6 | 2780.0 | 493.4 |
| | 2500 | 0.01 | 20000000 | 0 | 2030.0 | 512.4 | 3353.3 | 1404.4 | 2264.7 | 1185.4 |
| | | 0.05 | 20000000 | 0 | 2353.0 | 272.6 | 4115.0 | 1796.2 | 3833.7 | 502.7 |
| | | 0.1 | 20000000 | 0 | 1798.0 | 301.0 | 2350.0 | 1311.8 | 2284.7 | 696.6 |
| | | 0.2 | 20000000 | 0 | 3234.3 | 951.0 | 2072.7 | 695.8 | 3856.0 | 1951.3 |
| | | 0.4 | 20000000 | 0 | 2099.7 | 223.2 | 4429.3 | 1724.0 | 2849.3 | 461.1 |
| | | 0.8 | 20000000 | 0 | 2325.3 | 832.8 | 3014.3 | 758.9 | 3046.3 | 592.8 |
| | 3500 | 0.01 | 20000000 | 0 | 2396.3 | 1405.5 | 3955.0 | 1674.8 | 2475.7 | 545.3 |
| | | 0.05 | 20000000 | 0 | 2425.7 | 1046.6 | 3316.7 | 1439.5 | 2402.0 | 712.1 |
| | | 0.1 | 20000000 | 0 | 2050.0 | 179.9 | 2809.3 | 990.0 | 1568.0 | 717.8 |
| | | 0.2 | 20000000 | 0 | 2884.3 | 640.6 | 3457.3 | 2759.8 | 2381.7 | 948.9 |
| | | 0.4 | 20000000 | 0 | 1806.0 | 340.4 | 2576.7 | 896.7 | 2018.3 | 1410.0 |
| | | 0.8 | 20000000 | 0 | 2237.3 | 425.7 | 2184.7 | 641.9 | 3957.3 | 1103.6 |
| **Study case W2** 35M reads Length-weighted bias omitted | 1500 | 0.01 | 35000000 | 0 | 5727.3 | 4121.2 | 4075.3 | 1252.5 | 5042.7 | 379.9 |
| | | 0.05 | 35000000 | 0 | 3668.7 | 534.0 | 4029.0 | 1358.1 | 7460.7 | 6266.0 |
| | | 0.1 | 35000000 | 0 | 3390.0 | 1594.6 | 5918.0 | 3148.9 | 4773.0 | 531.1 |
| | | 0.2 | 35000000 | 0 | 4356.7 | 1146.1 | 4662.7 | 2078.9 | 4511.3 | 1009.4 |
| | | 0.4 | 35000000 | 0 | 4004.0 | 1084.6 | 4736.0 | 2104.3 | 6101.3 | 1819.6 |
| | | 0.8 | 35000000 | 0 | 4753.7 | 1696.5 | 4632.7 | 1564.6 | 3395.3 | 297.6 |
| | 2500 | 0.01 | 35000000 | 0 | 5530.0 | 2018.8 | 6097.7 | 2666.0 | 5858.0 | 1128.0 |
| | | 0.05 | 35000000 | 0 | 3726.7 | 1071.5 | 4736.3 | 1145.6 | 4566.7 | 1565.6 |
| | | 0.1 | 35000000 | 0 | 6178.0 | 1663.8 | 5543.3 | 1161.4 | 3704.7 | 433.7 |
| | | 0.2 | 35000000 | 0 | 6547.7 | 1064.0 | 3586.3 | 1572.6 | 3161.0 | 1487.0 |
| | | 0.4 | 35000000 | 0 | 3964.3 | 976.0 | 4440.0 | 1839.2 | 4954.7 | 1162.0 |
| | | 0.8 | 35000000 | 0 | 4229.3 | 1805.2 | 6121.0 | 808.9 | 4482.7 | 979.2 |
| | 3500 | 0.01 | 35000000 | 0 | 4266.3 | 1555.7 | 5605.7 | 1867.7 | 3819.0 | 525.5 |
| | | 0.05 | 35000000 | 0 | 3884.7 | 906.6 | 3830.0 | 1409.6 | 4241.7 | 1223.4 |
| | | 0.1 | 35000000 | 0 | 5113.0 | 436.9 | 4519.0 | 1135.9 | 5109.3 | 1787.3 |
| | | 0.2 | 35000000 | 0 | 4128.7 | 1893.5 | 4623.0 | 726.3 | 5275.7 | 1383.6 |
| | | 0.4 | 35000000 | 0 | 5122.3 | 3108.2 | 4633.0 | 1469.8 | 5468.0 | 1354.5 |
| | | 0.8 | 35000000 | 0 | 3824.0 | 997.5 | 3872.0 | 766.9 | 3973.0 | 2095.7 |
| **Study case W3** 35M reads Length-weighted: medium | 1500 | 0.01 | 35000000 | 0 | 2451.7 | 286.7 | 2581.0 | 317.5 | 2319.3 | 364.0 |
| | | 0.05 | 35000000 | 0 | 1967.0 | 75.5 | 2622.0 | 595.0 | 2648.0 | 336.1 |
| | | 0.1 | 35000000 | 0 | 2087.3 | 584.6 | 2555.7 | 222.3 | 2757.3 | 620.1 |
| | | 0.2 | 35000000 | 0 | 3054.3 | 930.1 | 2067.7 | 457.7 | 2266.7 | 848.8 |
| | | 0.4 | 35000000 | 0 | 2727.7 | 1058.7 | 2306.0 | 420.8 | 2980.7 | 617.3 |
| | | 0.8 | 35000000 | 0 | 2526.0 | 1012.0 | 2647.3 | 659.3 | 2581.3 | 1067.7 |
| | 2500 | 0.01 | 35000000 | 0 | 3287.3 | 1518.8 | 2682.3 | 1060.9 | 2149.3 | 449.7 |
| | | 0.05 | 35000000 | 0 | 2812.0 | 380.5 | 2379.0 | 963.0 | 2361.0 | 186.7 |
| | | 0.1 | 35000000 | 0 | 2576.3 | 405.0 | 2486.7 | 717.7 | 2922.0 | 1006.1 |
| | | 0.2 | 35000000 | 0 | 2995.3 | 1158.3 | 2790.7 | 1289.4 | 2830.7 | 646.4 |
| | | 0.4 | 35000000 | 0 | 2432.7 | 720.1 | 2898.3 | 600.3 | 2502.7 | 1233.6 |
| | | 0.8 | 35000000 | 0 | 2531.3 | 1268.6 | 2909.0 | 1015.3 | 2881.7 | 1277.5 |
| | 3500 | 0.01 | 35000000 | 0 | 2521.3 | 506.0 | 1969.7 | 277.1 | 2543.0 | 1031.4 |
| | | 0.05 | 35000000 | 0 | 3823.3 | 511.5 | 1928.3 | 222.8 | 2314.3 | 61.8 |
| | | 0.1 | 35000000 | 0 | 2064.7 | 1038.1 | 2252.7 | 1060.0 | 2092.0 | 322.1 |
| | | 0.2 | 35000000 | 0 | 2005.0 | 272.7 | 3616.7 | 1563.0 | 2837.7 | 939.6 |
| | | 0.4 | 35000000 | 0 | 2083.3 | 117.8 | 2283.7 | 1026.6 | 3059.7 | 1294.7 |
| | | 0.8 | 35000000 | 0 | 2522.7 | 758.1 | 2177.0 | 24.3 | 2514.7 | 558.5 |

| Non/mutated sample in study cases | Chr1 copy number | Fraction mutated copies | Raw reads | | Unmapped reads | |
|---|---|---|---|---|---|---|
| | | | Mean | SD | Mean | SD |
| **Study case W1** | 1500 | 0 | 20000000 | 0 | 2308.7 | 119.3 |
| | 2500 | 0 | 20000000 | 0 | 3387.0 | 2095.7 |
| | 3500 | 0 | 20000000 | 0 | 2009.3 | 605.1 |
| **Study case W2** | 1500 | 0 | 35000000 | 0 | 4050.7 | 903.6 |
| | 2500 | 0 | 35000000 | 0 | 4439.7 | 1990.9 |
| | 3500 | 0 | 35000000 | 0 | 4440.7 | 1683.9 |
| **Study case W3** | 1500 | 0 | 35000000 | 0 | 2090.0 | 446.1 |
| | 2500 | 0 | 35000000 | 0 | 2425.3 | 1036.0 |
| | 3500 | 0 | 35000000 | 0 | 2215.0 | 936.5 |

Table S9. Overview of the percentage of covered and not covered exons in study case W1. The table shows mean and SD of exon percentage covered with more than 10x, 50x, 100x, 200x and 300x in all samples, as well as the percentage of uncovered exons grouped by the lengths 1-50 bp, > 50 bp.

| | Percentage of mutated positions | Chr1 copy number | Fraction of mutated copies | Percentage of exons covered with more than | | | | | | | | | | Percentage of exons not covered | | | | | |
|---|---|---|---|---|---|---|---|---|---|---|---|---|---|---|---|---|---|---|---|
| | | | | 10x | | 50x | | 100x | | 200x | | 300x | | Lengths | | | | | |
| | | | | | | | | | | | | | | All | | 1 - 50 bp | | > 50 bp | |
| | | | | Mean | SD | Mean | SD | Mean | SD | Mean | SD | Mean | SD | Mean | SD | Mean | SD | Mean | SD |
| Study vase W1 - 20M No length-weight sequencing bias | Non-mutated | 1500 | 0 | 99.91 | 0.05 | 95.29 | 6.61 | 36.87 | 49.51 | 0.02 | 0.03 | 0.00 | 0.00 | 0.09 | 0.05 | 0.09 | 0.05 | 0 | 0 |
| | | 2500 | 0 | 99.95 | 0.07 | 94.84 | 8.65 | 60.04 | 52.15 | 0.09 | 0.12 | 0.01 | 0.01 | 0.05 | 0.07 | 0.05 | 0.07 | 0 | 0 |
| | | 3500 | 0 | 99.99 | 0.01 | 99.89 | 0.09 | 95.10 | 6.19 | 2.02 | 3.26 | 0.01 | 0.01 | 0.01 | 0.01 | 0.01 | 0.01 | 0 | 0 |
| | 0.5% | 1500 | 0.01 | 100.0 | 0.0 | 97.6 | 3.9 | 64.9 | 55.0 | 2.7 | 4.7 | 0.0 | 0.0 | 0.0 | 0.0 | 0.0 | 0.0 | 0.0 | 0.0 |
| | | | 0.05 | 100.0 | 0.0 | 99.9 | 0.1 | 96.6 | 4.8 | 6.1 | 6.7 | 0.0 | 0.0 | 0.0 | 0.0 | 0.0 | 0.0 | 0.0 | 0.0 |
| | | | 0.1 | 100.0 | 0.0 | 99.9 | 0.0 | 98.3 | 1.3 | 1.8 | 2.2 | 0.0 | 0.0 | 0.0 | 0.0 | 0.0 | 0.0 | 0.0 | 0.0 |
| | | | 0.2 | 100.0 | 0.0 | 100.0 | 0.0 | 99.3 | 0.1 | 6.6 | 4.5 | 0.0 | 0.0 | 0.0 | 0.0 | 0.0 | 0.0 | 0.0 | 0.0 |
| | | | 0.4 | 100.0 | 0.0 | 99.5 | 0.5 | 60.2 | 35.1 | 4.0 | 6.9 | 0.0 | 0.0 | 0.0 | 0.0 | 0.0 | 0.0 | 0.0 | 0.0 |
| | | | 0.8 | 100.0 | 0.1 | 97.1 | 4.9 | 66.1 | 56.1 | 0.5 | 0.6 | 0.0 | 0.0 | 0.0 | 0.1 | 0.0 | 0.1 | 0.0 | 0.0 |
| | | 2500 | 0.01 | 100.0 | 0.0 | 99.7 | 0.2 | 81.5 | 20.9 | 0.0 | 0.0 | 0.0 | 0.0 | 0.0 | 0.0 | 0.0 | 0.0 | 0.0 | 0.0 |
| | | | 0.05 | 99.9 | 0.2 | 74.0 | 44.8 | 54.2 | 50.3 | 4.3 | 7.4 | 0.0 | 0.0 | 0.1 | 0.1 | 0.1 | 0.1 | 0.0 | 0.0 |
| | | | 0.1 | 100.0 | 0.0 | 99.8 | 0.1 | 84.9 | 12.3 | 0.1 | 0.1 | 0.0 | 0.0 | 0.0 | 0.0 | 0.0 | 0.0 | 0.0 | 0.0 |
| | | | 0.2 | 100.0 | 0.0 | 99.4 | 0.7 | 67.3 | 40.7 | 0.0 | 0.0 | 0.0 | 0.0 | 0.0 | 0.0 | 0.0 | 0.0 | 0.0 | 0.0 |
| | | | 0.4 | 99.8 | 0.4 | 69.2 | 53.4 | 66.2 | 57.4 | 4.4 | 3.9 | 0.0 | 0.0 | 0.1 | 0.2 | 0.1 | 0.2 | 0.0 | 0.0 |
| | | | 0.8 | 100.0 | 0.0 | 99.9 | 0.1 | 96.1 | 5.7 | 5.1 | 4.8 | 0.0 | 0.0 | 0.0 | 0.0 | 0.0 | 0.0 | 0.0 | 0.0 |
| | | 3500 | 0.01 | 100.0 | 0.0 | 99.9 | 0.1 | 94.7 | 8.0 | 3.7 | 3.3 | 0.0 | 0.0 | 0.0 | 0.0 | 0.0 | 0.0 | 0.0 | 0.0 |
| | | | 0.05 | 100.0 | 0.0 | 99.4 | 0.8 | 63.3 | 40.5 | 0.1 | 0.1 | 0.0 | 0.0 | 0.0 | 0.0 | 0.0 | 0.0 | 0.0 | 0.0 |
| | | | 0.1 | 100.0 | 0.0 | 99.9 | 0.1 | 94.7 | 7.8 | 4.2 | 5.4 | 0.0 | 0.0 | 0.0 | 0.0 | 0.0 | 0.0 | 0.0 | 0.0 |
| | | | 0.2 | 100.0 | 0.0 | 99.9 | 0.0 | 96.8 | 2.7 | 1.9 | 3.1 | 0.0 | 0.0 | 0.0 | 0.0 | 0.0 | 0.0 | 0.0 | 0.0 |
| | | | 0.4 | 100.0 | 0.0 | 99.7 | 0.2 | 82.6 | 16.0 | 2.4 | 4.1 | 0.0 | 0.0 | 0.0 | 0.0 | 0.0 | 0.0 | 0.0 | 0.0 |
| | | | 0.8 | 100.0 | 0.0 | 99.9 | 0.2 | 93.7 | 9.9 | 4.2 | 4.0 | 0.0 | 0.0 | 0.0 | 0.0 | 0.0 | 0.0 | 0.0 | 0.0 |
| | 1% | 1500 | 0.01 | 100.0 | 0.0 | 99.8 | 0.2 | 86.5 | 11.1 | 0.7 | 1.1 | 0.0 | 0.0 | 0.0 | 0.0 | 0.0 | 0.0 | 0.0 | 0.0 |
| | | | 0.05 | 100.0 | 0.0 | 99.6 | 0.6 | 74.9 | 38.8 | 0.2 | 0.1 | 0.0 | 0.0 | 0.0 | 0.0 | 0.0 | 0.0 | 0.0 | 0.0 |
| | | | 0.1 | 100.0 | 0.0 | 99.8 | 0.2 | 83.9 | 21.3 | 0.1 | 0.2 | 0.0 | 0.0 | 0.0 | 0.0 | 0.0 | 0.0 | 0.0 | 0.0 |
| | | | 0.2 | 99.9 | 0.1 | 87.4 | 21.3 | 52.0 | 49.8 | 2.4 | 4.2 | 0.0 | 0.0 | 0.1 | 0.1 | 0.1 | 0.1 | 0.0 | 0.0 |
| | | | 0.4 | 100.0 | 0.0 | 99.9 | 0.1 | 92.0 | 10.4 | 1.8 | 3.0 | 0.0 | 0.0 | 0.0 | 0.0 | 0.0 | 0.0 | 0.0 | 0.0 |
| | | | 0.8 | 100.0 | 0.0 | 99.8 | 0.1 | 84.7 | 13.9 | 0.5 | 0.9 | 0.0 | 0.0 | 0.0 | 0.0 | 0.0 | 0.0 | 0.0 | 0.0 |
| | | 2500 | 0.01 | 99.9 | 0.2 | 72.7 | 47.2 | 65.7 | 56.9 | 3.8 | 6.5 | 0.0 | 0.0 | 0.1 | 0.1 | 0.1 | 0.1 | 0.0 | 0.0 |
| | | | 0.05 | 100.0 | 0.0 | 100.0 | 0.0 | 98.3 | 1.9 | 8.8 | 7.6 | 0.0 | 0.0 | 0.0 | 0.0 | 0.0 | 0.0 | 0.0 | 0.0 |
| | | | 0.1 | 100.0 | 0.0 | 99.9 | 0.0 | 99.1 | 0.6 | 6.7 | 5.9 | 0.0 | 0.0 | 0.0 | 0.0 | 0.0 | 0.0 | 0.0 | 0.0 |
| | | | 0.2 | 100.0 | 0.0 | 99.9 | 0.1 | 95.5 | 4.0 | 3.0 | 5.1 | 0.0 | 0.0 | 0.0 | 0.0 | 0.0 | 0.0 | 0.0 | 0.0 |
| | | | 0.4 | 100.0 | 0.0 | 99.9 | 0.2 | 92.2 | 12.6 | 6.8 | 6.2 | 0.0 | 0.0 | 0.0 | 0.0 | 0.0 | 0.0 | 0.0 | 0.0 |
| | | | 0.8 | 100.0 | 0.0 | 99.0 | 1.4 | 63.0 | 47.5 | 0.0 | 0.0 | 0.0 | 0.0 | 0.0 | 0.0 | 0.0 | 0.0 | 0.0 | 0.0 |
| | | 3500 | 0.01 | 100.0 | 0.0 | 98.9 | 1.5 | 56.8 | 43.0 | 0.0 | 0.0 | 0.0 | 0.0 | 0.0 | 0.0 | 0.0 | 0.0 | 0.0 | 0.0 |
| | | | 0.05 | 100.0 | 0.0 | 100.0 | 0.0 | 99.3 | 0.1 | 4.5 | 1.2 | 0.0 | 0.0 | 0.0 | 0.0 | 0.0 | 0.0 | 0.0 | 0.0 |
| | | | 0.1 | 100.0 | 0.0 | 99.9 | 0.1 | 93.9 | 6.5 | 0.3 | 0.4 | 0.0 | 0.0 | 0.0 | 0.0 | 0.0 | 0.0 | 0.0 | 0.0 |
| | | | 0.2 | 100.0 | 0.0 | 99.9 | 0.1 | 95.8 | 5.1 | 2.1 | 3.4 | 0.0 | 0.0 | 0.0 | 0.0 | 0.0 | 0.0 | 0.0 | 0.0 |
| | | | 0.4 | 100.0 | 0.0 | 99.7 | 0.2 | 72.8 | 23.0 | 0.9 | 1.5 | 0.0 | 0.0 | 0.0 | 0.0 | 0.0 | 0.0 | 0.0 | 0.0 |
| | | | 0.8 | 100.0 | 0.0 | 99.7 | 0.3 | 78.5 | 21.8 | 3.8 | 6.6 | 0.0 | 0.0 | 0.0 | 0.0 | 0.0 | 0.0 | 0.0 | 0.0 |
| | 5% | 1500 | 0.01 | 100.0 | 0.0 | 99.9 | 0.1 | 94.4 | 7.0 | 1.5 | 2.4 | 0.0 | 0.0 | 0.0 | 0.0 | 0.0 | 0.0 | 0.0 | 0.0 |
| | | | 0.05 | 100.0 | 0.0 | 99.4 | 1.0 | 71.1 | 48.7 | 3.0 | 3.7 | 0.0 | 0.0 | 0.0 | 0.0 | 0.0 | 0.0 | 0.0 | 0.0 |
| | | | 0.1 | 100.0 | 0.0 | 99.8 | 0.1 | 85.7 | 14.2 | 0.0 | 0.0 | 0.0 | 0.0 | 0.0 | 0.0 | 0.0 | 0.0 | 0.0 | 0.0 |
| | | | 0.2 | 100.0 | 0.0 | 98.4 | 2.1 | 48.5 | 47.1 | 0.1 | 0.2 | 0.0 | 0.0 | 0.0 | 0.0 | 0.0 | 0.0 | 0.0 | 0.0 |
| | | | 0.4 | 100.0 | 0.0 | 99.2 | 1.0 | 62.6 | 43.9 | 0.1 | 0.1 | 0.0 | 0.0 | 0.0 | 0.0 | 0.0 | 0.0 | 0.0 | 0.0 |
| | | | 0.8 | 100.0 | 0.0 | 99.8 | 0.1 | 91.7 | 9.3 | 0.1 | 0.1 | 0.0 | 0.0 | 0.0 | 0.0 | 0.0 | 0.0 | 0.0 | 0.0 |
| | | 2500 | 0.01 | 99.9 | 0.0 | 97.3 | 4.1 | 53.3 | 45.7 | 0.0 | 0.0 | 0.0 | 0.0 | 0.1 | 0.0 | 0.1 | 0.0 | 0.0 | 0.0 |
| | | | 0.05 | 100.0 | 0.0 | 99.8 | 0.2 | 89.0 | 17.4 | 1.8 | 2.1 | 0.0 | 0.0 | 0.0 | 0.0 | 0.0 | 0.0 | 0.0 | 0.0 |
| | | | 0.1 | 100.0 | 0.0 | 99.7 | 0.2 | 82.2 | 14.5 | 0.0 | 0.0 | 0.0 | 0.0 | 0.0 | 0.0 | 0.0 | 0.0 | 0.0 | 0.0 |
| | | | 0.2 | 100.0 | 0.0 | 99.9 | 0.1 | 97.5 | 1.8 | 3.0 | 5.0 | 0.0 | 0.0 | 0.0 | 0.0 | 0.0 | 0.0 | 0.0 | 0.0 |
| | | | 0.4 | 100.0 | 0.0 | 99.4 | 0.8 | 72.2 | 44.3 | 1.1 | 1.8 | 0.0 | 0.0 | 0.0 | 0.0 | 0.0 | 0.0 | 0.0 | 0.0 |
| | | | 0.8 | 100.0 | 0.0 | 99.5 | 0.5 | 70.1 | 37.0 | 0.0 | 0.0 | 0.0 | 0.0 | 0.0 | 0.0 | 0.0 | 0.0 | 0.0 | 0.0 |
| | | 3500 | 0.01 | 100.0 | 0.0 | 99.9 | 0.1 | 94.6 | 6.4 | 2.8 | 4.8 | 0.0 | 0.0 | 0.0 | 0.0 | 0.0 | 0.0 | 0.0 | 0.0 |
| | | | 0.05 | 100.0 | 0.0 | 99.9 | 0.1 | 95.9 | 6.1 | 6.5 | 6.5 | 0.0 | 0.0 | 0.0 | 0.0 | 0.0 | 0.0 | 0.0 | 0.0 |
| | | | 0.1 | 100.0 | 0.0 | 99.9 | 0.1 | 96.8 | 4.4 | 6.1 | 5.7 | 0.0 | 0.0 | 0.0 | 0.0 | 0.0 | 0.0 | 0.0 | 0.0 |
| | | | 0.2 | 99.9 | 0.1 | 87.0 | 22.2 | 59.5 | 52.4 | 1.3 | 2.2 | 0.0 | 0.0 | 0.1 | 0.1 | 0.1 | 0.1 | 0.0 | 0.0 |
| | | | 0.4 | 100.0 | 0.0 | 99.3 | 0.9 | 60.9 | 42.6 | 1.1 | 1.9 | 0.0 | 0.0 | 0.0 | 0.0 | 0.0 | 0.0 | 0.0 | 0.0 |
| | | | 0.8 | 100.0 | 0.0 | 99.9 | 0.1 | 95.8 | 4.6 | 1.5 | 2.3 | 0.0 | 0.0 | 0.0 | 0.0 | 0.0 | 0.0 | 0.0 | 0.0 |

Table S10. Overview of the percentage of covered and not covered exons in study case W2. The table shows mean and SD of exon percentage covered with more than 10x, 50x, 100x, 200x and 300x in all samples, as well as the percentage of uncovered exons grouped by the lengths 1-50 bp, > 50 bp.

| | Percentage of mutated positions | Chr1 copy number | Fraction of mutated copies | Percentage of exons covered with more then | | | | | | | | | | Percentage of exons not covered | | | | | |
|---|---|---|---|---|---|---|---|---|---|---|---|---|---|---|---|---|---|---|---|
| | | | | *10x* | | *50x* | | *100x* | | *200x* | | *300x* | | *Lengths* | | | | | |
| | | | | | | | | | | | | | | *All* | | *1 - 50 bp* | | *>50 bp* | |
| | | | | Mean | SD | Mean | SD | Mean | SD | Mean | SD | Mean | SD | Mean | SD | Mean | SD | Mean | SD |
| **Study vase W2 - 35M No length-weight sequencing bias** | **Non-mutated** | 1500 | 0 | 100 | 0 | 100 | 0 | 99.99 | 0.00 | 99.62 | 0.01 | 92.48 | 0.27 | 0 | 0 | 0 | 0 | 0 | 0 |
| | | 2500 | 0 | 100 | 0 | 100 | 0 | 99.99 | 0.00 | 99.61 | 0.00 | 92.93 | 0.18 | 0 | 0 | 0 | 0 | 0 | 0 |
| | | 3500 | 0 | 100 | 0 | 100 | 0 | 99.99 | 0.00 | 99.62 | 0.01 | 92.52 | 0.49 | 0 | 0 | 0 | 0 | 0 | 0 |
| | **0.5%** | 1500 | 0.01 | 100.0 | 0.0 | 100.0 | 0.0 | 100.0 | 0.0 | 99.6 | 0.0 | 92.6 | 0.2 | 0.0 | 0.0 | 0.0 | 0.0 | 0.0 | 0.0 |
| | | | 0.05 | 100.0 | 0.0 | 100.0 | 0.0 | 100.0 | 0.0 | 99.6 | 0.0 | 92.5 | 0.3 | 0.0 | 0.0 | 0.0 | 0.0 | 0.0 | 0.0 |
| | | | 0.1 | 100.0 | 0.0 | 100.0 | 0.0 | 100.0 | 0.0 | 99.6 | 0.0 | 93.2 | 0.2 | 0.0 | 0.0 | 0.0 | 0.0 | 0.0 | 0.0 |
| | | | 0.2 | 100.0 | 0.0 | 100.0 | 0.0 | 100.0 | 0.0 | 99.6 | 0.0 | 92.6 | 0.2 | 0.0 | 0.0 | 0.0 | 0.0 | 0.0 | 0.0 |
| | | | 0.4 | 100.0 | 0.0 | 100.0 | 0.0 | 100.0 | 0.0 | 99.6 | 0.0 | 92.6 | 0.3 | 0.0 | 0.0 | 0.0 | 0.0 | 0.0 | 0.0 |
| | | | 0.8 | 100.0 | 0.0 | 100.0 | 0.0 | 100.0 | 0.0 | 99.6 | 0.0 | 92.9 | 0.1 | 0.0 | 0.0 | 0.0 | 0.0 | 0.0 | 0.0 |
| | | 2500 | 0.01 | 100.0 | 0.0 | 100.0 | 0.0 | 100.0 | 0.0 | 99.6 | 0.0 | 92.7 | 0.1 | 0.0 | 0.0 | 0.0 | 0.0 | 0.0 | 0.0 |
| | | | 0.05 | 100.0 | 0.0 | 100.0 | 0.0 | 100.0 | 0.0 | 99.6 | 0.0 | 92.7 | 0.6 | 0.0 | 0.0 | 0.0 | 0.0 | 0.0 | 0.0 |
| | | | 0.1 | 100.0 | 0.0 | 100.0 | 0.0 | 100.0 | 0.0 | 99.6 | 0.0 | 92.8 | 0.5 | 0.0 | 0.0 | 0.0 | 0.0 | 0.0 | 0.0 |
| | | | 0.2 | 100.0 | 0.0 | 100.0 | 0.0 | 100.0 | 0.0 | 99.6 | 0.0 | 92.5 | 0.2 | 0.0 | 0.0 | 0.0 | 0.0 | 0.0 | 0.0 |
| | | | 0.4 | 100.0 | 0.0 | 100.0 | 0.0 | 100.0 | 0.0 | 99.6 | 0.0 | 92.6 | 0.4 | 0.0 | 0.0 | 0.0 | 0.0 | 0.0 | 0.0 |
| | | | 0.8 | 100.0 | 0.0 | 100.0 | 0.0 | 100.0 | 0.0 | 99.6 | 0.0 | 92.6 | 0.4 | 0.0 | 0.0 | 0.0 | 0.0 | 0.0 | 0.0 |
| | | 3500 | 0.01 | 100.0 | 0.0 | 100.0 | 0.0 | 100.0 | 0.0 | 99.6 | 0.0 | 92.9 | 0.5 | 0.0 | 0.0 | 0.0 | 0.0 | 0.0 | 0.0 |
| | | | 0.05 | 100.0 | 0.0 | 100.0 | 0.0 | 100.0 | 0.0 | 99.6 | 0.0 | 93.1 | 0.1 | 0.0 | 0.0 | 0.0 | 0.0 | 0.0 | 0.0 |
| | | | 0.1 | 100.0 | 0.0 | 100.0 | 0.0 | 100.0 | 0.0 | 99.6 | 0.0 | 93.0 | 0.2 | 0.0 | 0.0 | 0.0 | 0.0 | 0.0 | 0.0 |
| | | | 0.2 | 100.0 | 0.0 | 100.0 | 0.0 | 100.0 | 0.0 | 99.6 | 0.0 | 93.1 | 0.4 | 0.0 | 0.0 | 0.0 | 0.0 | 0.0 | 0.0 |
| | | | 0.4 | 100.0 | 0.0 | 100.0 | 0.0 | 100.0 | 0.0 | 99.6 | 0.0 | 93.0 | 0.5 | 0.0 | 0.0 | 0.0 | 0.0 | 0.0 | 0.0 |
| | | | 0.8 | 100.0 | 0.0 | 100.0 | 0.0 | 100.0 | 0.0 | 99.6 | 0.0 | 92.8 | 0.4 | 0.0 | 0.0 | 0.0 | 0.0 | 0.0 | 0.0 |
| | **1%** | 1500 | 0.01 | 100.0 | 0.0 | 100.0 | 0.0 | 100.0 | 0.0 | 99.6 | 0.0 | 92.5 | 0.2 | 0.0 | 0.0 | 0.0 | 0.0 | 0.0 | 0.0 |
| | | | 0.05 | 100.0 | 0.0 | 100.0 | 0.0 | 100.0 | 0.0 | 99.6 | 0.0 | 92.5 | 0.3 | 0.0 | 0.0 | 0.0 | 0.0 | 0.0 | 0.0 |
| | | | 0.1 | 100.0 | 0.0 | 100.0 | 0.0 | 100.0 | 0.0 | 99.6 | 0.0 | 92.6 | 0.4 | 0.0 | 0.0 | 0.0 | 0.0 | 0.0 | 0.0 |
| | | | 0.2 | 100.0 | 0.0 | 100.0 | 0.0 | 100.0 | 0.0 | 99.6 | 0.0 | 92.5 | 0.3 | 0.0 | 0.0 | 0.0 | 0.0 | 0.0 | 0.0 |
| | | | 0.4 | 100.0 | 0.0 | 100.0 | 0.0 | 100.0 | 0.0 | 99.6 | 0.0 | 92.3 | 0.4 | 0.0 | 0.0 | 0.0 | 0.0 | 0.0 | 0.0 |
| | | | 0.8 | 100.0 | 0.0 | 100.0 | 0.0 | 100.0 | 0.0 | 99.6 | 0.0 | 92.6 | 0.1 | 0.0 | 0.0 | 0.0 | 0.0 | 0.0 | 0.0 |
| | | 2500 | 0.01 | 100.0 | 0.0 | 100.0 | 0.0 | 100.0 | 0.0 | 99.6 | 0.0 | 92.8 | 0.5 | 0.0 | 0.0 | 0.0 | 0.0 | 0.0 | 0.0 |
| | | | 0.05 | 100.0 | 0.0 | 100.0 | 0.0 | 100.0 | 0.0 | 99.6 | 0.0 | 92.7 | 0.1 | 0.0 | 0.0 | 0.0 | 0.0 | 0.0 | 0.0 |
| | | | 0.1 | 100.0 | 0.0 | 100.0 | 0.0 | 100.0 | 0.0 | 99.6 | 0.0 | 92.7 | 0.3 | 0.0 | 0.0 | 0.0 | 0.0 | 0.0 | 0.0 |
| | | | 0.2 | 100.0 | 0.0 | 100.0 | 0.0 | 100.0 | 0.0 | 99.6 | 0.0 | 93.3 | 0.4 | 0.0 | 0.0 | 0.0 | 0.0 | 0.0 | 0.0 |
| | | | 0.4 | 100.0 | 0.0 | 100.0 | 0.0 | 100.0 | 0.0 | 99.6 | 0.0 | 92.7 | 0.9 | 0.0 | 0.0 | 0.0 | 0.0 | 0.0 | 0.0 |
| | | | 0.8 | 100.0 | 0.0 | 100.0 | 0.0 | 100.0 | 0.0 | 99.6 | 0.0 | 92.0 | 0.2 | 0.0 | 0.0 | 0.0 | 0.0 | 0.0 | 0.0 |
| | | 3500 | 0.01 | 100.0 | 0.0 | 100.0 | 0.0 | 100.0 | 0.0 | 99.6 | 0.0 | 92.8 | 0.4 | 0.0 | 0.0 | 0.0 | 0.0 | 0.0 | 0.0 |
| | | | 0.05 | 100.0 | 0.0 | 100.0 | 0.0 | 100.0 | 0.0 | 99.6 | 0.0 | 93.0 | 0.3 | 0.0 | 0.0 | 0.0 | 0.0 | 0.0 | 0.0 |
| | | | 0.1 | 100.0 | 0.0 | 100.0 | 0.0 | 100.0 | 0.0 | 99.6 | 0.0 | 92.8 | 0.1 | 0.0 | 0.0 | 0.0 | 0.0 | 0.0 | 0.0 |
| | | | 0.2 | 100.0 | 0.0 | 100.0 | 0.0 | 100.0 | 0.0 | 99.6 | 0.0 | 93.1 | 0.4 | 0.0 | 0.0 | 0.0 | 0.0 | 0.0 | 0.0 |
| | | | 0.4 | 100.0 | 0.0 | 100.0 | 0.0 | 100.0 | 0.0 | 99.6 | 0.0 | 93.2 | 0.3 | 0.0 | 0.0 | 0.0 | 0.0 | 0.0 | 0.0 |
| | | | 0.8 | 100.0 | 0.0 | 100.0 | 0.0 | 100.0 | 0.0 | 99.6 | 0.0 | 92.7 | 0.9 | 0.0 | 0.0 | 0.0 | 0.0 | 0.0 | 0.0 |
| | **5%** | 1500 | 0.01 | 100.0 | 0.0 | 100.0 | 0.0 | 100.0 | 0.0 | 99.6 | 0.0 | 92.6 | 0.4 | 0.0 | 0.0 | 0.0 | 0.0 | 0.0 | 0.0 |
| | | | 0.05 | 100.0 | 0.0 | 100.0 | 0.0 | 100.0 | 0.0 | 99.6 | 0.0 | 92.1 | 0.2 | 0.0 | 0.0 | 0.0 | 0.0 | 0.0 | 0.0 |
| | | | 0.1 | 100.0 | 0.0 | 100.0 | 0.0 | 100.0 | 0.0 | 99.6 | 0.0 | 92.8 | 0.2 | 0.0 | 0.0 | 0.0 | 0.0 | 0.0 | 0.0 |
| | | | 0.2 | 100.0 | 0.0 | 100.0 | 0.0 | 100.0 | 0.0 | 99.6 | 0.0 | 92.5 | 0.1 | 0.0 | 0.0 | 0.0 | 0.0 | 0.0 | 0.0 |
| | | | 0.4 | 100.0 | 0.0 | 100.0 | 0.0 | 100.0 | 0.0 | 99.6 | 0.0 | 92.8 | 0.5 | 0.0 | 0.0 | 0.0 | 0.0 | 0.0 | 0.0 |
| | | | 0.8 | 100.0 | 0.0 | 100.0 | 0.0 | 100.0 | 0.0 | 99.6 | 0.0 | 92.8 | 0.3 | 0.0 | 0.0 | 0.0 | 0.0 | 0.0 | 0.0 |
| | | 2500 | 0.01 | 100.0 | 0.0 | 100.0 | 0.0 | 100.0 | 0.0 | 99.6 | 0.0 | 93.1 | 0.2 | 0.0 | 0.0 | 0.0 | 0.0 | 0.0 | 0.0 |
| | | | 0.05 | 100.0 | 0.0 | 100.0 | 0.0 | 100.0 | 0.0 | 99.6 | 0.0 | 93.1 | 0.4 | 0.0 | 0.0 | 0.0 | 0.0 | 0.0 | 0.0 |
| | | | 0.1 | 100.0 | 0.0 | 100.0 | 0.0 | 100.0 | 0.0 | 99.6 | 0.0 | 92.8 | 0.3 | 0.0 | 0.0 | 0.0 | 0.0 | 0.0 | 0.0 |
| | | | 0.2 | 100.0 | 0.0 | 100.0 | 0.0 | 100.0 | 0.0 | 99.6 | 0.0 | 92.6 | 0.5 | 0.0 | 0.0 | 0.0 | 0.0 | 0.0 | 0.0 |
| | | | 0.4 | 100.0 | 0.0 | 100.0 | 0.0 | 100.0 | 0.0 | 99.6 | 0.0 | 92.4 | 0.4 | 0.0 | 0.0 | 0.0 | 0.0 | 0.0 | 0.0 |
| | | | 0.8 | 100.0 | 0.0 | 100.0 | 0.0 | 100.0 | 0.0 | 99.6 | 0.0 | 92.5 | 0.2 | 0.0 | 0.0 | 0.0 | 0.0 | 0.0 | 0.0 |
| | | 3500 | 0.01 | 100.0 | 0.0 | 100.0 | 0.0 | 100.0 | 0.0 | 99.6 | 0.0 | 93.1 | 0.5 | 0.0 | 0.0 | 0.0 | 0.0 | 0.0 | 0.0 |
| | | | 0.05 | 100.0 | 0.0 | 100.0 | 0.0 | 100.0 | 0.0 | 99.6 | 0.0 | 92.7 | 0.5 | 0.0 | 0.0 | 0.0 | 0.0 | 0.0 | 0.0 |
| | | | 0.1 | 100.0 | 0.0 | 100.0 | 0.0 | 100.0 | 0.0 | 99.6 | 0.0 | 93.0 | 0.5 | 0.0 | 0.0 | 0.0 | 0.0 | 0.0 | 0.0 |
| | | | 0.2 | 100.0 | 0.0 | 100.0 | 0.0 | 100.0 | 0.0 | 99.6 | 0.0 | 92.4 | 0.2 | 0.0 | 0.0 | 0.0 | 0.0 | 0.0 | 0.0 |
| | | | 0.4 | 100.0 | 0.0 | 100.0 | 0.0 | 100.0 | 0.0 | 99.6 | 0.0 | 92.8 | 0.1 | 0.0 | 0.0 | 0.0 | 0.0 | 0.0 | 0.0 |
| | | | 0.8 | 100.0 | 0.0 | 100.0 | 0.0 | 100.0 | 0.0 | 99.6 | 0.0 | 92.9 | 0.3 | 0.0 | 0.0 | 0.0 | 0.0 | 0.0 | 0.0 |

Table S11. Overview of the percentage of covered and not covered exons in study case W3. The table shows mean and SD of exon percentage covered with more than 10x, 50x, 100x, 200x and 300x in all samples, as well as the percentage of uncovered exons grouped by the lengths 1-50 bp, > 50 bp.

| | Percentage of mutated positions | Chr1 copy number | Fraction of mutated copies | Percentage of exons covered with more then | | | | | | | | | | Percentage of exons not covered | | | | | |
|---|---|---|---|---|---|---|---|---|---|---|---|---|---|---|---|---|---|---|---|
| | | | | | | | | | | | | | | *Lengths* | | | | | |
| | | | | *10x* | | *50x* | | *100x* | | *200x* | | *300x* | | *All* | | *1 - 50 bp* | | *>50 bp* | |
| | | | | Mean | SD | Mean | SD | Mean | SD | Mean | SD | Mean | SD | Mean | SD | Mean | SD | Mean | SD |
| **Study vase W3- 35M Length-weight sequencing bias: Median** | **Non-mutated** | 1500 | 0 | 100 | 0 | 100 | 0 | 100 | 0 | 99.90 | 0.00 | 99.20 | 0.04 | 0 | 0 | 0 | 0 | 0 | 0 |
| | | 2500 | 0 | 100 | 0 | 100 | 0 | 100 | 0 | 99.90 | 0.01 | 99.20 | 0.02 | 0 | 0 | 0 | 0 | 0 | 0 |
| | | 3500 | 0 | 100 | 0 | 100 | 0 | 100 | 0 | 99.90 | 0.01 | 99.20 | 0.03 | 0 | 0 | 0 | 0 | 0 | 0 |
| | **0.5%** | 1500 | 0.01 | 100.0 | 0.0 | 100.0 | 0.0 | 100.0 | 0.0 | 99.9 | 0.0 | 99.1 | 0.0 | 0.0 | 0.0 | 0.0 | 0.0 | 0.0 | 0.0 |
| | | | 0.05 | 100.0 | 0.0 | 100.0 | 0.0 | 100.0 | 0.0 | 99.9 | 0.0 | 99.2 | 0.0 | 0.0 | 0.0 | 0.0 | 0.0 | 0.0 | 0.0 |
| | | | 0.1 | 100.0 | 0.0 | 100.0 | 0.0 | 100.0 | 0.0 | 99.9 | 0.0 | 99.2 | 0.0 | 0.0 | 0.0 | 0.0 | 0.0 | 0.0 | 0.0 |
| | | | 0.2 | 100.0 | 0.0 | 100.0 | 0.0 | 100.0 | 0.0 | 99.9 | 0.0 | 99.2 | 0.0 | 0.0 | 0.0 | 0.0 | 0.0 | 0.0 | 0.0 |
| | | | 0.4 | 100.0 | 0.0 | 100.0 | 0.0 | 100.0 | 0.0 | 99.9 | 0.0 | 99.2 | 0.0 | 0.0 | 0.0 | 0.0 | 0.0 | 0.0 | 0.0 |
| | | | 0.8 | 100.0 | 0.0 | 100.0 | 0.0 | 100.0 | 0.0 | 99.9 | 0.0 | 99.2 | 0.0 | 0.0 | 0.0 | 0.0 | 0.0 | 0.0 | 0.0 |
| | | 2500 | 0.01 | 100.0 | 0.0 | 100.0 | 0.0 | 100.0 | 0.0 | 99.9 | 0.0 | 99.2 | 0.0 | 0.0 | 0.0 | 0.0 | 0.0 | 0.0 | 0.0 |
| | | | 0.05 | 100.0 | 0.0 | 100.0 | 0.0 | 100.0 | 0.0 | 99.9 | 0.0 | 99.2 | 0.0 | 0.0 | 0.0 | 0.0 | 0.0 | 0.0 | 0.0 |
| | | | 0.1 | 100.0 | 0.0 | 100.0 | 0.0 | 100.0 | 0.0 | 99.9 | 0.0 | 99.2 | 0.0 | 0.0 | 0.0 | 0.0 | 0.0 | 0.0 | 0.0 |
| | | | 0.2 | 100.0 | 0.0 | 100.0 | 0.0 | 100.0 | 0.0 | 99.9 | 0.0 | 99.2 | 0.0 | 0.0 | 0.0 | 0.0 | 0.0 | 0.0 | 0.0 |
| | | | 0.4 | 100.0 | 0.0 | 100.0 | 0.0 | 100.0 | 0.0 | 99.9 | 0.0 | 99.2 | 0.0 | 0.0 | 0.0 | 0.0 | 0.0 | 0.0 | 0.0 |
| | | | 0.8 | 100.0 | 0.0 | 100.0 | 0.0 | 100.0 | 0.0 | 99.9 | 0.0 | 99.2 | 0.0 | 0.0 | 0.0 | 0.0 | 0.0 | 0.0 | 0.0 |
| | | 3500 | 0.01 | 100.0 | 0.0 | 100.0 | 0.0 | 100.0 | 0.0 | 99.9 | 0.0 | 99.2 | 0.0 | 0.0 | 0.0 | 0.0 | 0.0 | 0.0 | 0.0 |
| | | | 0.05 | 100.0 | 0.0 | 100.0 | 0.0 | 100.0 | 0.0 | 99.9 | 0.0 | 99.2 | 0.0 | 0.0 | 0.0 | 0.0 | 0.0 | 0.0 | 0.0 |
| | | | 0.1 | 100.0 | 0.0 | 100.0 | 0.0 | 100.0 | 0.0 | 99.9 | 0.0 | 99.2 | 0.0 | 0.0 | 0.0 | 0.0 | 0.0 | 0.0 | 0.0 |
| | | | 0.2 | 100.0 | 0.0 | 100.0 | 0.0 | 100.0 | 0.0 | 99.9 | 0.0 | 99.2 | 0.0 | 0.0 | 0.0 | 0.0 | 0.0 | 0.0 | 0.0 |
| | | | 0.4 | 100.0 | 0.0 | 100.0 | 0.0 | 100.0 | 0.0 | 99.9 | 0.0 | 99.2 | 0.0 | 0.0 | 0.0 | 0.0 | 0.0 | 0.0 | 0.0 |
| | | | 0.8 | 100.0 | 0.0 | 100.0 | 0.0 | 100.0 | 0.0 | 99.9 | 0.0 | 99.3 | 0.0 | 0.0 | 0.0 | 0.0 | 0.0 | 0.0 | 0.0 |
| | **1%** | 1500 | 0.01 | 100.0 | 0.0 | 100.0 | 0.0 | 100.0 | 0.0 | 99.9 | 0.0 | 99.2 | 0.0 | 0.0 | 0.0 | 0.0 | 0.0 | 0.0 | 0.0 |
| | | | 0.05 | 100.0 | 0.0 | 100.0 | 0.0 | 100.0 | 0.0 | 99.9 | 0.0 | 99.2 | 0.0 | 0.0 | 0.0 | 0.0 | 0.0 | 0.0 | 0.0 |
| | | | 0.1 | 100.0 | 0.0 | 100.0 | 0.0 | 100.0 | 0.0 | 99.9 | 0.0 | 99.2 | 0.0 | 0.0 | 0.0 | 0.0 | 0.0 | 0.0 | 0.0 |
| | | | 0.2 | 100.0 | 0.0 | 100.0 | 0.0 | 100.0 | 0.0 | 99.9 | 0.0 | 99.2 | 0.0 | 0.0 | 0.0 | 0.0 | 0.0 | 0.0 | 0.0 |
| | | | 0.4 | 100.0 | 0.0 | 100.0 | 0.0 | 100.0 | 0.0 | 99.9 | 0.0 | 99.1 | 0.0 | 0.0 | 0.0 | 0.0 | 0.0 | 0.0 | 0.0 |
| | | | 0.8 | 100.0 | 0.0 | 100.0 | 0.0 | 100.0 | 0.0 | 99.9 | 0.0 | 99.2 | 0.1 | 0.0 | 0.0 | 0.0 | 0.0 | 0.0 | 0.0 |
| | | 2500 | 0.01 | 100.0 | 0.0 | 100.0 | 0.0 | 100.0 | 0.0 | 99.9 | 0.0 | 99.2 | 0.0 | 0.0 | 0.0 | 0.0 | 0.0 | 0.0 | 0.0 |
| | | | 0.05 | 100.0 | 0.0 | 100.0 | 0.0 | 100.0 | 0.0 | 99.9 | 0.0 | 99.2 | 0.0 | 0.0 | 0.0 | 0.0 | 0.0 | 0.0 | 0.0 |
| | | | 0.1 | 100.0 | 0.0 | 100.0 | 0.0 | 100.0 | 0.0 | 99.9 | 0.0 | 99.2 | 0.0 | 0.0 | 0.0 | 0.0 | 0.0 | 0.0 | 0.0 |
| | | | 0.2 | 100.0 | 0.0 | 100.0 | 0.0 | 100.0 | 0.0 | 99.9 | 0.0 | 99.2 | 0.0 | 0.0 | 0.0 | 0.0 | 0.0 | 0.0 | 0.0 |
| | | | 0.4 | 100.0 | 0.0 | 100.0 | 0.0 | 100.0 | 0.0 | 99.9 | 0.0 | 99.2 | 0.0 | 0.0 | 0.0 | 0.0 | 0.0 | 0.0 | 0.0 |
| | | | 0.8 | 100.0 | 0.0 | 100.0 | 0.0 | 100.0 | 0.0 | 99.9 | 0.0 | 99.2 | 0.0 | 0.0 | 0.0 | 0.0 | 0.0 | 0.0 | 0.0 |
| | | 3500 | 0.01 | 100.0 | 0.0 | 100.0 | 0.0 | 100.0 | 0.0 | 99.9 | 0.0 | 99.2 | 0.1 | 0.0 | 0.0 | 0.0 | 0.0 | 0.0 | 0.0 |
| | | | 0.05 | 100.0 | 0.0 | 100.0 | 0.0 | 100.0 | 0.0 | 99.9 | 0.0 | 99.2 | 0.0 | 0.0 | 0.0 | 0.0 | 0.0 | 0.0 | 0.0 |
| | | | 0.1 | 100.0 | 0.0 | 100.0 | 0.0 | 100.0 | 0.0 | 99.9 | 0.0 | 99.2 | 0.0 | 0.0 | 0.0 | 0.0 | 0.0 | 0.0 | 0.0 |
| | | | 0.2 | 100.0 | 0.0 | 100.0 | 0.0 | 100.0 | 0.0 | 99.9 | 0.0 | 99.2 | 0.0 | 0.0 | 0.0 | 0.0 | 0.0 | 0.0 | 0.0 |
| | | | 0.4 | 100.0 | 0.0 | 100.0 | 0.0 | 100.0 | 0.0 | 99.9 | 0.0 | 99.2 | 0.0 | 0.0 | 0.0 | 0.0 | 0.0 | 0.0 | 0.0 |
| | | | 0.8 | 100.0 | 0.0 | 100.0 | 0.0 | 100.0 | 0.0 | 99.9 | 0.0 | 99.2 | 0.1 | 0.0 | 0.0 | 0.0 | 0.0 | 0.0 | 0.0 |
| | **5%** | 1500 | 0.01 | 100 | 0 | 100.0 | 0.0 | 100.0 | 0.0 | 99.9 | 0.0 | 99.2 | 0.0 | 0.0 | 0.0 | 0.0 | 0.0 | 0.0 | 0.0 |
| | | | 0.05 | 100 | 0 | 100.0 | 0.0 | 100.0 | 0.0 | 99.9 | 0.0 | 99.2 | 0.0 | 0.0 | 0.0 | 0.0 | 0.0 | 0.0 | 0.0 |
| | | | 0.1 | 100 | 0 | 100.0 | 0.0 | 100.0 | 0.0 | 99.9 | 0.0 | 99.1 | 0.0 | 0.0 | 0.0 | 0.0 | 0.0 | 0.0 | 0.0 |
| | | | 0.2 | 100 | 0 | 100.0 | 0.0 | 100.0 | 0.0 | 99.9 | 0.0 | 99.2 | 0.0 | 0.0 | 0.0 | 0.0 | 0.0 | 0.0 | 0.0 |
| | | | 0.4 | 100 | 0 | 100.0 | 0.0 | 100.0 | 0.0 | 99.9 | 0.0 | 99.2 | 0.0 | 0.0 | 0.0 | 0.0 | 0.0 | 0.0 | 0.0 |
| | | | 0.8 | 100 | 0 | 100.0 | 0.0 | 100.0 | 0.0 | 99.9 | 0.0 | 99.1 | 0.0 | 0.0 | 0.0 | 0.0 | 0.0 | 0.0 | 0.0 |
| | | 2500 | 0.01 | 100 | 0 | 100.0 | 0.0 | 100.0 | 0.0 | 99.9 | 0.0 | 99.2 | 0.0 | 0.0 | 0.0 | 0.0 | 0.0 | 0.0 | 0.0 |
| | | | 0.05 | 100 | 0 | 100.0 | 0.0 | 100.0 | 0.0 | 99.9 | 0.0 | 99.2 | 0.0 | 0.0 | 0.0 | 0.0 | 0.0 | 0.0 | 0.0 |
| | | | 0.1 | 100 | 0 | 100.0 | 0.0 | 100.0 | 0.0 | 99.9 | 0.0 | 99.2 | 0.0 | 0.0 | 0.0 | 0.0 | 0.0 | 0.0 | 0.0 |
| | | | 0.2 | 100 | 0 | 100.0 | 0.0 | 100.0 | 0.0 | 99.9 | 0.0 | 99.2 | 0.0 | 0.0 | 0.0 | 0.0 | 0.0 | 0.0 | 0.0 |
| | | | 0.4 | 100 | 0 | 100.0 | 0.0 | 100.0 | 0.0 | 99.9 | 0.0 | 99.2 | 0.0 | 0.0 | 0.0 | 0.0 | 0.0 | 0.0 | 0.0 |
| | | | 0.8 | 100 | 0 | 100.0 | 0.0 | 100.0 | 0.0 | 99.9 | 0.0 | 99.2 | 0.0 | 0.0 | 0.0 | 0.0 | 0.0 | 0.0 | 0.0 |
| | | 3500 | 0.01 | 100 | 0 | 100.0 | 0.0 | 100.0 | 0.0 | 99.9 | 0.0 | 99.2 | 0.0 | 0.0 | 0.0 | 0.0 | 0.0 | 0.0 | 0.0 |
| | | | 0.05 | 100 | 0 | 100.0 | 0.0 | 100.0 | 0.0 | 99.9 | 0.0 | 99.2 | 0.0 | 0.0 | 0.0 | 0.0 | 0.0 | 0.0 | 0.0 |
| | | | 0.1 | 100 | 0 | 100.0 | 0.0 | 100.0 | 0.0 | 99.9 | 0.0 | 99.2 | 0.0 | 0.0 | 0.0 | 0.0 | 0.0 | 0.0 | 0.0 |
| | | | 0.2 | 100.0 | 0.0 | 100.0 | 0.0 | 99.9 | 0.0 | 99.2 | 0.0 | 0.0 | 0.0 | 0.0 | 0.0 | 0.0 | 0.0 | 0.0 | 0.0 |
| | | | 0.4 | 100.0 | 0.0 | 100.0 | 0.0 | 99.9 | 0.0 | 99.2 | 0.0 | 0.0 | 0.0 | 0.0 | 0.0 | 0.0 | 0.0 | 0.0 | 0.0 |
| | | | 0.8 | 100.0 | 0.0 | 100.0 | 0.0 | 100.0 | 0.0 | 99.9 | 0.0 | 99.4 | 0.0 | 0.0 | 0.0 | 0.0 | 0.0 | 0.0 | 0.0 |

Table S12. Overview of the inserted mutations in samples in study cases W1-3. The table exhibits the total number of unique mutations, mutation frequency in terms of the number of mutations per Mb DNA in an *in-silico* sample, and mean variant allele frequency (VAF).

| Percentage of mutated positions | Copy number | Fraction of mutated copies | Study case W1 20M No length-weighted bias | | | Study case W2 35M No length-weighted bias | | | Study case W3 35M Length-weighted bias: median | | |
|---|---|---|---|---|---|---|---|---|---|---|---|
| | | | Total number of unique mutation | Mutation frequency | Mean variant allele frequency | Total number of unique mutation | Mutation frequency | Mean variant allele frequency | Total number of unique mutation | Mutation frequency | Mean variant allele frequency |
| 0.5% | 1500 | 0.01 | 2307 | 0.05 | 0.004 | 2307 | 0.05 | 0.004 | 2306 | 0.04 | 0.004 |
| | | 0.05 | 2307 | 0.27 | 0.03 | 2307 | 0.24 | 0.03 | 2307 | 0.26 | 0.03 |
| | | 0.1 | 2307 | 0.47 | 0.04 | 2307 | 0.47 | 0.04 | 2307 | 0.41 | 0.04 |
| | | 0.2 | 2307 | 0.93 | 0.1 | 2307 | 0.92 | 0.1 | 2307 | 0.92 | 0.1 |
| | | 0.4 | 2307 | 1.84 | 0.2 | 2307 | 1.82 | 0.2 | 2307 | 1.76 | 0.2 |
| | | 0.8 | 2307 | 3.73 | 0.4 | 2307 | 3.71 | 0.4 | 2307 | 3.69 | 0.4 |
| | 2500 | 0.01 | 2307 | 0.05 | 0.005 | 2307 | 0.05 | 0.005 | 2307 | 0.04 | 0.005 |
| | | 0.05 | 2307 | 0.21 | 0.02 | 2307 | 0.22 | 0.02 | 2307 | 0.21 | 0.02 |
| | | 0.1 | 2307 | 0.50 | 0.05 | 2307 | 0.44 | 0.05 | 2307 | 0.45 | 0.05 |
| | | 0.2 | 2307 | 0.93 | 0.1 | 2307 | 0.91 | 0.1 | 2307 | 0.91 | 0.1 |
| | | 0.4 | 2307 | 1.91 | 0.2 | 2307 | 1.84 | 0.2 | 2308 | 1.85 | 0.2 |
| | | 0.8 | 2307 | 3.68 | 0.4 | 2307 | 3.68 | 0.4 | 2307 | 3.74 | 0.4 |
| | 3500 | 0.01 | 2307 | 0.05 | 0.005 | 2307 | 0.05 | 0.005 | 2307 | 0.05 | 0.005 |
| | | 0.05 | 2307 | 0.22 | 0.02 | 2307 | 0.23 | 0.02 | 2307 | 0.23 | 0.02 |
| | | 0.1 | 2307 | 0.48 | 0.05 | 2307 | 0.47 | 0.05 | 2307 | 0.46 | 0.05 |
| | | 0.2 | 2307 | 0.96 | 0.1 | 2307 | 0.94 | 0.1 | 2307 | 0.94 | 0.1 |
| | | 0.4 | 2307 | 1.84 | 0.2 | 2307 | 1.83 | 0.2 | 2307 | 1.87 | 0.2 |
| | | 0.8 | 2308 | 3.70 | 0.4 | 2307 | 3.73 | 0.4 | 2307 | 3.66 | 0.4 |
| 1% | 1500 | 0.01 | 4613 | 0.10 | 0.005 | 4614 | 0.10 | 0.005 | 4613 | 0.09 | 0.005 |
| | | 0.05 | 4614 | 0.44 | 0.03 | 4614 | 0.45 | 0.03 | 4614 | 0.51 | 0.03 |
| | | 0.1 | 4614 | 0.95 | 0.05 | 4615 | 0.94 | 0.05 | 4614 | 0.88 | 0.05 |
| | | 0.2 | 4614 | 1.85 | 0.1 | 4614 | 1.88 | 0.1 | 4614 | 1.78 | 0.1 |
| | | 0.4 | 4614 | 3.84 | 0.2 | 4614 | 3.62 | 0.2 | 4614 | 3.71 | 0.2 |
| | | 0.8 | 4614 | 7.53 | 0.4 | 4614 | 7.67 | 0.4 | 4614 | 7.35 | 0.4 |
| | 2500 | 0.01 | 4614 | 0.11 | 0.005 | 4614 | 0.09 | 0.005 | 4614 | 0.10 | 0.005 |
| | | 0.05 | 4614 | 0.47 | 0.03 | 4614 | 0.47 | 0.03 | 4614 | 0.51 | 0.03 |
| | | 0.1 | 4614 | 0.93 | 0.05 | 4614 | 0.90 | 0.05 | 4614 | 0.92 | 0.05 |
| | | 0.2 | 4615 | 1.88 | 0.1 | 4614 | 1.80 | 0.1 | 4614 | 1.86 | 0.1 |
| | | 0.4 | 4614 | 3.71 | 0.2 | 4615 | 3.78 | 0.2 | 4614 | 3.74 | 0.2 |
| | | 0.8 | 4614 | 7.55 | 0.4 | 4614 | 7.46 | 0.4 | 4614 | 7.40 | 0.4 |
| | 3500 | 0.01 | 4614 | 0.07 | 0.005 | 4614 | 0.10 | 0.005 | 4614 | 0.08 | 0.005 |
| | | 0.05 | 4614 | 0.46 | 0.03 | 4614 | 0.45 | 0.03 | 4614 | 0.47 | 0.03 |
| | | 0.1 | 4614 | 0.95 | 0.05 | 4614 | 0.94 | 0.05 | 4615 | 0.90 | 0.05 |
| | | 0.2 | 4614 | 1.89 | 0.1 | 4614 | 1.87 | 0.1 | 4614 | 1.80 | 0.1 |
| | | 0.4 | 4614 | 3.68 | 0.2 | 4614 | 3.71 | 0.2 | 4614 | 3.68 | 0.2 |
| | | 0.8 | 4615 | 7.40 | 0.4 | 4615 | 7.46 | 0.4 | 4614 | 7.46 | 0.4 |
| 5% | 1500 | 0.01 | 23069 | 0.41 | 0.006 | 23069 | 0.52 | 0.006 | 23067 | 0.52 | 0.006 |
| | | 0.05 | 23068 | 2.04 | 0.03 | 23069 | 2.46 | 0.03 | 23069 | 2.42 | 0.03 |
| | | 0.1 | 23068 | 4.56 | 0.05 | 23068 | 4.60 | 0.05 | 23068 | 4.81 | 0.05 |
| | | 0.2 | 23069 | 9.38 | 0.1 | 23069 | 9.17 | 0.1 | 23068 | 9.50 | 0.1 |
| | | 0.4 | 23068 | 19.03 | 0.2 | 23068 | 18.22 | 0.2 | 23068 | 18.94 | 0.2 |
| | | 0.8 | 23068 | 35.90 | 0.4 | 23068 | 37.75 | 0.4 | 23069 | 37.05 | 0.4 |
| | 2500 | 0.01 | 23068 | 0.39 | 0.004 | 23068 | 0.47 | 0.004 | 23068 | 0.42 | 0.004 |
| | | 0.05 | 23070 | 2.09 | 0.03 | 23068 | 2.42 | 0.03 | 23068 | 2.41 | 0.03 |
| | | 0.1 | 23069 | 4.82 | 0.05 | 23070 | 4.69 | 0.05 | 23070 | 4.78 | 0.05 |
| | | 0.2 | 23068 | 8.96 | 0.1 | 23069 | 9.65 | 0.1 | 23069 | 9.17 | 0.1 |
| | | 0.4 | 23068 | 18.63 | 0.2 | 23068 | 18.62 | 0.2 | 23068 | 18.42 | 0.2 |
| | | 0.8 | 23070 | 37.09 | 0.4 | 23070 | 36.88 | 0.4 | 23068 | 36.90 | 0.4 |
| | 3500 | 0.01 | 23068 | 0.43 | 0.005 | 23068 | 0.50 | 0.005 | 23069 | 0.46 | 0.005 |
| | | 0.05 | 23070 | 2.38 | 0.03 | 23068 | 2.32 | 0.03 | 23070 | 2.44 | 0.03 |
| | | 0.1 | 23069 | 4.74 | 0.05 | 23069 | 4.73 | 0.05 | 23069 | 4.69 | 0.05 |
| | | 0.2 | 23068 | 9.41 | 0.1 | 23068 | 8.85 | 0.1 | 23068 | 9.33 | 0.1 |
| | | 0.4 | 23069 | 18.55 | 0.2 | 23068 | 18.00 | 0.2 | 23068 | 18.52 | 0.2 |
| | | 0.8 | 23068 | 37.08 | 0.4 | 23070 | 36.57 | 0.4 | 23068 | 37.19 | 0.4 |

Table S13. Overview of the mean precision and recall, and mean and SD of true positives, false positives, and false negatives in samples from study case W1.

| Percentage of mutated positions | Copy number | Fraction of mutated genomes | Mean precision | Mean recall | TP | | FP | | FN | |
|---|---|---|---|---|---|---|---|---|---|---|
| | | | | | Mean | SD | Mean | SD | Mean | SD |
| 0.5% | 1500 | 0.01 | 0.9 | 0.0 | 45.7 | 18.0 | 4.7 | 0.6 | 2261.3 | 18.0 |
| | | 0.05 | 1.0 | 0.5 | 1050.0 | 223.3 | 22.0 | 18.4 | 1257.0 | 223.3 |
| | | 0.1 | 0.9 | 0.7 | 1718.0 | 12.2 | 142.0 | 116.9 | 589.0 | 12.2 |
| | | 0.2 | 1.0 | 0.9 | 1972.0 | 5.3 | 7.3 | 4.0 | 335.0 | 5.3 |
| | | 0.4 | 1.0 | 0.9 | 1986.7 | 0.6 | 24.0 | 8.7 | 320.3 | 0.6 |
| | | 0.8 | 1.0 | 0.8 | 1951.0 | 0.0 | 12.0 | 2.6 | 356.0 | 0.0 |
| | 2500 | 0.01 | 0.8 | 0.0 | 43.7 | 22.2 | 9.0 | 7.9 | 2263.3 | 22.2 |
| | | 0.05 | 1.0 | 0.4 | 891.3 | 48.2 | 18.0 | 11.8 | 1415.7 | 48.2 |
| | | 0.1 | 1.0 | 0.8 | 1757.3 | 42.7 | 94.0 | 82.2 | 549.7 | 42.7 |
| | | 0.2 | 1.0 | 0.8 | 1959.3 | 3.2 | 11.0 | 2.6 | 347.7 | 3.2 |
| | | 0.4 | 1.0 | 0.9 | 1962.7 | 2.9 | 16.7 | 4.9 | 344.3 | 2.9 |
| | | 0.8 | 1.0 | 0.9 | 2001.0 | 2.6 | 6.7 | 4.6 | 306.0 | 2.6 |
| | 3500 | 0.01 | 1.0 | 0.0 | 45.7 | 30.6 | 2.0 | 3.5 | 2261.3 | 30.6 |
| | | 0.05 | 1.0 | 0.5 | 1068.0 | 206.4 | 19.7 | 6.8 | 1239.0 | 206.4 |
| | | 0.1 | 1.0 | 0.8 | 1779.3 | 28.2 | 24.7 | 6.5 | 527.7 | 28.2 |
| | | 0.2 | 1.0 | 0.8 | 1933.0 | 3.0 | 3.7 | 3.2 | 374.0 | 3.0 |
| | | 0.4 | 1.0 | 0.9 | 1979.7 | 4.7 | 12.3 | 9.7 | 327.3 | 4.7 |
| | | 0.8 | 1.0 | 0.8 | 1956.3 | 3.8 | 4.0 | 1.7 | 351.7 | 3.8 |
| 1 % | 1500 | 0.01 | 0.9 | 0.0 | 103.3 | 52.6 | 15.7 | 19.3 | 4509.7 | 52.6 |
| | | 0.05 | 1.0 | 0.4 | 1719.0 | 47.4 | 28.0 | 21.8 | 2895.0 | 47.4 |
| | | 0.1 | 1.0 | 0.8 | 3548.0 | 29.8 | 43.0 | 19.5 | 1066.0 | 29.8 |
| | | 0.2 | 1.0 | 0.8 | 3912.0 | 16.7 | 27.3 | 20.6 | 702.0 | 16.7 |
| | | 0.4 | 1.0 | 0.8 | 3902.0 | 3.5 | 21.7 | 24.6 | 712.0 | 3.5 |
| | | 0.8 | 1.0 | 0.9 | 3957.7 | 6.1 | 12.7 | 10.0 | 656.3 | 6.1 |
| | 2500 | 0.01 | 0.9 | 0.0 | 59.3 | 24.0 | 10.3 | 16.2 | 4554.7 | 24.0 |
| | | 0.05 | 0.9 | 0.4 | 1678.7 | 273.0 | 91.7 | 107.3 | 2935.3 | 273.0 |
| | | 0.1 | 1.0 | 0.8 | 3529.0 | 65.6 | 24.0 | 19.1 | 1085.0 | 65.6 |
| | | 0.2 | 1.0 | 0.8 | 3860.0 | 7.5 | 13.0 | 4.6 | 755.0 | 7.5 |
| | | 0.4 | 1.0 | 0.8 | 3921.3 | 8.3 | 21.3 | 8.1 | 692.7 | 8.3 |
| | | 0.8 | 1.0 | 0.8 | 3916.0 | 9.6 | 9.0 | 6.1 | 698.0 | 9.6 |
| | 3500 | 0.01 | 0.8 | 0.0 | 23.3 | 14.0 | 5.7 | 4.5 | 4590.7 | 14.0 |
| | | 0.05 | 1.0 | 0.3 | 1407.3 | 340.8 | 27.3 | 6.5 | 3206.7 | 340.8 |
| | | 0.1 | 1.0 | 0.8 | 3491.7 | 53.6 | 127.7 | 161.3 | 1122.3 | 53.6 |
| | | 0.2 | 1.0 | 0.8 | 3866.0 | 4.6 | 7.7 | 1.5 | 748.0 | 4.6 |
| | | 0.4 | 1.0 | 0.8 | 3918.0 | 10.0 | 14.0 | 5.3 | 696.0 | 10.0 |
| | | 0.8 | 1.0 | 0.9 | 3954.3 | 10.0 | 4.0 | 1.7 | 660.7 | 10.0 |
| 5 % | 1500 | 0.01 | 1.0 | 0.0 | 367.7 | 170.1 | 17.0 | 11.1 | 22701.3 | 170.1 |
| | | 0.05 | 1.0 | 0.3 | 6904.0 | 266.1 | 51.0 | 31.8 | 16164.0 | 266.1 |
| | | 0.1 | 1.0 | 0.6 | 13153.7 | 357.5 | 89.7 | 83.9 | 9914.3 | 357.5 |
| | | 0.2 | 1.0 | 0.6 | 14737.7 | 72.9 | 105.0 | 89.7 | 8331.3 | 72.9 |
| | | 0.4 | 1.0 | 0.6 | 14963.0 | 28.4 | 25.7 | 18.0 | 8105.0 | 28.4 |
| | | 0.8 | 1.0 | 0.6 | 14894.7 | 36.7 | 10.0 | 3.5 | 8173.3 | 36.7 |
| | 2500 | 0.01 | 0.9 | 0.0 | 293.0 | 41.9 | 17.7 | 9.0 | 22775.0 | 41.9 |
| | | 0.05 | 1.0 | 0.3 | 7030.0 | 1305.8 | 177.3 | 129.4 | 16040.0 | 1305.8 |
| | | 0.1 | 1.0 | 0.6 | 13562.3 | 304.0 | 150.3 | 161.0 | 9506.7 | 304.0 |
| | | 0.2 | 1.0 | 0.6 | 14691.7 | 25.7 | 72.0 | 71.9 | 8376.3 | 25.7 |
| | | 0.4 | 1.0 | 0.6 | 14980.7 | 13.6 | 20.0 | 12.1 | 8087.3 | 13.6 |
| | | 0.8 | 1.0 | 0.6 | 14786.0 | 40.8 | 13.7 | 2.1 | 8284.0 | 40.8 |
| | 3500 | 0.01 | 0.8 | 0.0 | 434.3 | 219.1 | 186.3 | 298.6 | 22633.7 | 219.1 |
| | | 0.05 | 1.0 | 0.3 | 7989.0 | 1248.4 | 242.3 | 386.8 | 15081.0 | 1248.4 |
| | | 0.1 | 1.0 | 0.6 | 13801.3 | 100.6 | 168.0 | 248.9 | 9267.7 | 100.6 |
| | | 0.2 | 1.0 | 0.6 | 14842.3 | 13.8 | 11.0 | 5.6 | 8225.7 | 13.8 |
| | | 0.4 | 1.0 | 0.6 | 14886.0 | 26.9 | 8.0 | 6.1 | 8183.0 | 26.9 |
| | | 0.8 | 1.0 | 0.6 | 14896.0 | 61.1 | 12.0 | 3.5 | 8172.0 | 61.1 |

Table S14. Overview of the mean precision and recall, and mean and SD of true positives, false positives, and false negatives in samples from study case W2.

| Percentage of mutated positions | Copy number | Fraction of mutated genomes | Mean precision | Mean recall | TP | | FP | | FN | |
|---|---|---|---|---|---|---|---|---|---|---|
| | | | | | Mean | SD | Mean | SD | Mean | SD |
| 0.5% | 1500 | 0.01 | 0.9 | 0.0 | 61.7 | 51.4 | 3.7 | 0.6 | 2245.3 | 51.4 |
| | | 0.05 | 1.0 | 0.5 | 1108.3 | 75.1 | 9.3 | 4.9 | 1198.7 | 75.1 |
| | | 0.1 | 1.0 | 0.8 | 1939.7 | 18.0 | 82.0 | 93.6 | 367.3 | 18.0 |
| | | 0.2 | 1.0 | 0.9 | 1996.0 | 2.6 | 14.7 | 8.1 | 311.0 | 2.6 |
| | | 0.4 | 1.0 | 0.9 | 1974.7 | 3.8 | 8.7 | 3.8 | 332.3 | 3.8 |
| | | 0.8 | 1.0 | 0.9 | 1974.0 | 2.6 | 6.0 | 7.8 | 333.0 | 2.6 |
| | 2500 | 0.01 | 0.9 | 0.0 | 87.3 | 9.5 | 4.7 | 2.1 | 2219.7 | 9.5 |
| | | 0.05 | 0.9 | 0.6 | 1282.7 | 90.9 | 94.3 | 36.9 | 1024.3 | 90.9 |
| | | 0.1 | 1.0 | 0.8 | 1915.0 | 25.2 | 12.3 | 6.1 | 392.0 | 25.2 |
| | | 0.2 | 1.0 | 0.9 | 2005.0 | 7.0 | 7.7 | 5.0 | 302.0 | 7.0 |
| | | 0.4 | 1.0 | 0.9 | 1986.0 | 2.0 | 16.0 | 7.8 | 321.0 | 2.0 |
| | | 0.8 | 1.0 | 0.9 | 1990.3 | 2.9 | 4.7 | 0.6 | 316.7 | 2.9 |
| | 3500 | 0.01 | 1.0 | 0.0 | 97.7 | 14.4 | 1.3 | 1.5 | 2209.3 | 14.4 |
| | | 0.05 | 1.0 | 0.6 | 1269.7 | 142.4 | 13.0 | 3.6 | 1037.3 | 142.4 |
| | | 0.1 | 1.0 | 0.8 | 1892.0 | 25.1 | 11.3 | 6.4 | 415.0 | 25.1 |
| | | 0.2 | 1.0 | 0.9 | 1970.3 | 3.5 | 8.7 | 0.6 | 336.7 | 3.5 |
| | | 0.4 | 1.0 | 0.9 | 1973.7 | 2.1 | 11.7 | 1.5 | 333.3 | 2.1 |
| | | 0.8 | 1.0 | 0.9 | 2009.0 | 1.7 | 3.3 | 2.9 | 298.0 | 1.7 |
| 1 % | 1500 | 0.01 | 0.9 | 0.0 | 195.0 | 96.0 | 15.3 | 21.4 | 4419.0 | 96.0 |
| | | 0.05 | 0.9 | 0.5 | 2139.7 | 767.5 | 111.3 | 123.8 | 2474.3 | 767.5 |
| | | 0.1 | 1.0 | 0.8 | 3786.0 | 44.0 | 20.3 | 4.9 | 829.0 | 44.0 |
| | | 0.2 | 1.0 | 0.9 | 3922.3 | 6.0 | 48.3 | 32.7 | 691.7 | 6.0 |
| | | 0.4 | 1.0 | 0.9 | 3948.3 | 4.0 | 26.7 | 0.6 | 682.7 | 7.3 |
| | | 0.8 | 1.0 | 0.9 | 3960.3 | 1.2 | 7.7 | 9.1 | 653.7 | 1.2 |
| | 2500 | 0.01 | 0.9 | 0.1 | 249.0 | 72.5 | 13.0 | 12.2 | 4365.0 | 72.5 |
| | | 0.05 | 1.0 | 0.5 | 2419.3 | 172.1 | 38.3 | 18.0 | 2194.7 | 172.1 |
| | | 0.1 | 1.0 | 0.8 | 3727.0 | 12.5 | 23.3 | 23.2 | 887.0 | 12.5 |
| | | 0.2 | 1.0 | 0.9 | 3923.7 | 2.9 | 9.3 | 2.5 | 690.3 | 2.9 |
| | | 0.4 | 1.0 | 0.9 | 3949.0 | 4.6 | 10.7 | 6.7 | 666.0 | 4.6 |
| | | 0.8 | 1.0 | 0.9 | 3927.3 | 3.5 | 5.0 | 1.7 | 686.7 | 3.5 |
| | 3500 | 0.01 | 1.0 | 0.1 | 263.3 | 86.9 | 5.3 | 2.9 | 4350.7 | 86.9 |
| | | 0.05 | 1.0 | 0.6 | 2772.0 | 316.8 | 73.0 | 52.1 | 1842.0 | 316.8 |
| | | 0.1 | 1.0 | 0.8 | 3706.7 | 59.7 | 17.7 | 7.2 | 907.3 | 59.7 |
| | | 0.2 | 1.0 | 0.9 | 3963.3 | 3.8 | 10.7 | 2.5 | 650.7 | 3.8 |
| | | 0.4 | 1.0 | 0.8 | 3897.0 | 4.0 | 5.3 | 3.1 | 717.0 | 4.0 |
| | | 0.8 | 1.0 | 0.9 | 3955.0 | 13.0 | 18.3 | 8.5 | 660.0 | 13.0 |
| 5 % | 1500 | 0.01 | 1.0 | 0.0 | 793.0 | 196.0 | 7.3 | 2.3 | 22276.0 | 196.0 |
| | | 0.05 | 1.0 | 0.5 | 11050.7 | 1076.4 | 94.0 | 82.5 | 12018.3 | 1076.4 |
| | | 0.1 | 1.0 | 0.6 | 14457.7 | 124.0 | 77.0 | 51.5 | 8610.3 | 124.0 |
| | | 0.2 | 1.0 | 0.6 | 14866.3 | 67.0 | 38.3 | 33.5 | 8202.7 | 67.0 |
| | | 0.4 | 1.0 | 0.6 | 14907.3 | 22.5 | 9.3 | 4.0 | 8160.7 | 22.5 |
| | | 0.8 | 1.0 | 0.7 | 15120.3 | 48.4 | 13.0 | 4.6 | 7947.7 | 48.4 |
| | 2500 | 0.01 | 1.0 | 0.0 | 961.7 | 237.1 | 15.0 | 7.5 | 22106.3 | 237.1 |
| | | 0.05 | 1.0 | 0.5 | 11365.0 | 659.3 | 24.3 | 9.9 | 11703.0 | 659.3 |
| | | 0.1 | 1.0 | 0.6 | 14412.0 | 100.7 | 125.0 | 127.2 | 8658.0 | 100.7 |
| | | 0.2 | 1.0 | 0.6 | 14960.7 | 23.4 | 32.3 | 20.5 | 8108.3 | 23.4 |
| | | 0.4 | 1.0 | 0.6 | 14872.0 | 15.1 | 18.7 | 7.8 | 8196.0 | 15.1 |
| | | 0.8 | 1.0 | 0.7 | 15077.7 | 30.6 | 18.0 | 8.2 | 7992.3 | 30.6 |
| | 3500 | 0.01 | 1.0 | 0.0 | 882.7 | 137.3 | 4.3 | 2.5 | 22185.3 | 137.3 |
| | | 0.05 | 1.0 | 0.5 | 12562.7 | 382.0 | 30.3 | 19.7 | 10505.3 | 382.0 |
| | | 0.1 | 1.0 | 0.6 | 14602.0 | 42.0 | 22.7 | 2.9 | 8467.0 | 42.0 |
| | | 0.2 | 1.0 | 0.6 | 14884.0 | 64.9 | 35.3 | 28.7 | 8184.0 | 64.9 |
| | | 0.4 | 1.0 | 0.7 | 15037.0 | 89.2 | 22.0 | 10.6 | 8031.0 | 89.2 |
| | | 0.8 | 1.0 | 0.6 | 14904.0 | 30.4 | 21.0 | 9.5 | 8166.0 | 30.4 |

Table S15. Overview of the mean precision and recall, and mean and SD of true positives, false positives, and false negatives in samples from study case W3.

| Percentage of mutated positions | Copy number | Fraction of mutated genomes | Mean precision | Mean recall | TP | | FP | | FN | |
|---|---|---|---|---|---|---|---|---|---|---|
| | | | | | Mean | SD | Mean | SD | Mean | SD |
| 0.5% | 1500 | 0.01 | 0.9 | 0.0 | 85.7 | 40.5 | 10.3 | 5.1 | 2220.3 | 40.5 |
| | | 0.05 | 1.0 | 0.6 | 1443.7 | 134.9 | 16.0 | 10.4 | 863.3 | 134.9 |
| | | 0.1 | 1.0 | 0.8 | 1815.7 | 13.7 | 19.0 | 3.6 | 491.3 | 13.7 |
| | | 0.2 | 1.0 | 0.9 | 2003.7 | 5.0 | 18.0 | 1.7 | 303.3 | 5.0 |
| | | 0.4 | 1.0 | 0.9 | 1973.7 | 0.6 | 9.7 | 5.9 | 333.3 | 0.6 |
| | | 0.8 | 1.0 | 0.9 | 2011.3 | 0.6 | 8.7 | 1.5 | 295.7 | 0.6 |
| | 2500 | 0.01 | 1.0 | 0.1 | 129.3 | 8.3 | 6.7 | 0.6 | 2177.7 | 8.3 |
| | | 0.05 | 1.0 | 0.6 | 1269.0 | 99.8 | 45.7 | 59.3 | 1038.0 | 99.8 |
| | | 0.1 | 1.0 | 0.8 | 1906.3 | 13.4 | 20.7 | 9.3 | 400.7 | 13.4 |
| | | 0.2 | 1.0 | 0.9 | 1979.0 | 5.0 | 15.7 | 6.1 | 328.0 | 5.0 |
| | | 0.4 | 1.0 | 0.9 | 2011.3 | 2.5 | 9.0 | 3.5 | 296.7 | 2.5 |
| | | 0.8 | 1.0 | 0.9 | 1987.0 | 3.0 | 8.0 | 1.0 | 320.0 | 3.0 |
| | 3500 | 0.01 | 0.9 | 0.1 | 132.3 | 58.3 | 11.7 | 6.4 | 2174.7 | 58.3 |
| | | 0.05 | 1.0 | 0.6 | 1368.0 | 54.6 | 20.3 | 2.1 | 939.0 | 54.6 |
| | | 0.1 | 1.0 | 0.8 | 1946.0 | 8.7 | 24.7 | 11.6 | 361.0 | 8.7 |
| | | 0.2 | 1.0 | 0.9 | 1976.7 | 4.0 | 15.7 | 8.1 | 330.3 | 4.0 |
| | | 0.4 | 1.0 | 0.9 | 1991.7 | 6.7 | 7.7 | 2.3 | 315.3 | 6.7 |
| | | 0.8 | 1.0 | 0.9 | 1989.0 | 4.6 | 7.3 | 4.2 | 318.0 | 4.6 |
| 1 % | 1500 | 0.01 | 1.0 | 0.1 | 239.0 | 35.6 | 7.3 | 4.2 | 4374.0 | 35.6 |
| | | 0.05 | 1.0 | 0.6 | 2800.0 | 216.6 | 79.3 | 69.0 | 1814.0 | 216.6 |
| | | 0.1 | 1.0 | 0.8 | 3676.0 | 66.7 | 45.7 | 30.9 | 938.0 | 66.7 |
| | | 0.2 | 1.0 | 0.9 | 3924.0 | 7.8 | 17.3 | 1.5 | 690.0 | 7.8 |
| | | 0.4 | 1.0 | 0.9 | 3944.7 | 6.4 | 13.7 | 6.4 | 669.3 | 6.4 |
| | | 0.8 | 1.0 | 0.9 | 3948.3 | 4.0 | 11.3 | 2.1 | 665.7 | 4.0 |
| | 2500 | 0.01 | 1.0 | 0.0 | 175.3 | 64.7 | 7.7 | 4.0 | 4438.7 | 64.7 |
| | | 0.05 | 1.0 | 0.6 | 2955.3 | 211.2 | 60.7 | 37.8 | 1658.7 | 211.2 |
| | | 0.1 | 1.0 | 0.8 | 3770.3 | 29.8 | 19.0 | 7.5 | 843.7 | 29.8 |
| | | 0.2 | 1.0 | 0.8 | 3915.3 | 3.1 | 25.3 | 8.7 | 698.7 | 3.1 |
| | | 0.4 | 1.0 | 0.8 | 3907.0 | 5.2 | 14.7 | 2.9 | 707.0 | 5.2 |
| | | 0.8 | 1.0 | 0.9 | 3937.3 | 4.0 | 10.0 | 2.6 | 676.7 | 4.0 |
| | 3500 | 0.01 | 0.9 | 0.0 | 132.0 | 38.3 | 6.7 | 2.3 | 4482.0 | 38.3 |
| | | 0.05 | 1.0 | 0.6 | 2653.0 | 204.7 | 39.7 | 11.0 | 1961.0 | 204.7 |
| | | 0.1 | 1.0 | 0.8 | 3819.0 | 10.8 | 28.3 | 19.7 | 796.0 | 10.8 |
| | | 0.2 | 1.0 | 0.9 | 3965.3 | 12.7 | 54.7 | 75.7 | 648.7 | 12.7 |
| | | 0.4 | 1.0 | 0.8 | 3890.0 | 5.6 | 14.0 | 2.0 | 724.0 | 5.6 |
| | | 0.8 | 1.0 | 0.8 | 3915.7 | 9.5 | 14.0 | 2.0 | 698.3 | 9.5 |
| 5 % | 1500 | 0.01 | 1.0 | 0.0 | 896.0 | 104.8 | 9.7 | 5.5 | 22171.0 | 104.8 |
| | | 0.05 | 1.0 | 0.5 | 11337.3 | 124.7 | 66.3 | 42.0 | 11731.7 | 124.7 |
| | | 0.1 | 1.0 | 0.6 | 14552.7 | 85.6 | 38.3 | 8.5 | 8515.3 | 85.6 |
| | | 0.2 | 1.0 | 0.7 | 15134.7 | 27.1 | 24.7 | 6.7 | 7933.3 | 27.1 |
| | | 0.4 | 1.0 | 0.7 | 15239.0 | 6.1 | 25.7 | 3.1 | 7829.0 | 6.1 |
| | | 0.8 | 1.0 | 0.7 | 15085.0 | 37.7 | 16.7 | 3.8 | 7984.0 | 37.7 |
| | 2500 | 0.01 | 1.0 | 0.0 | 564.0 | 73.7 | 5.0 | 1.7 | 22504.0 | 73.7 |
| | | 0.05 | 1.0 | 0.5 | 12183.7 | 630.7 | 23.0 | 5.6 | 10884.3 | 630.7 |
| | | 0.1 | 1.0 | 0.6 | 14945.0 | 50.1 | 64.3 | 65.0 | 8125.0 | 50.1 |
| | | 0.2 | 1.0 | 0.7 | 15295.0 | 13.1 | 24.3 | 4.0 | 7774.0 | 13.1 |
| | | 0.4 | 1.0 | 0.7 | 15280.7 | 97.0 | 60.0 | 71.0 | 7787.3 | 97.0 |
| | | 0.8 | 1.0 | 0.7 | 15369.0 | 45.8 | 23.7 | 2.5 | 7699.0 | 45.8 |
| | 3500 | 0.01 | 1.0 | 0.0 | 748.7 | 193.8 | 3.3 | 1.5 | 22320.3 | 193.8 |
| | | 0.05 | 1.0 | 0.5 | 11977.3 | 544.8 | 38.0 | 15.7 | 11092.7 | 544.8 |
| | | 0.1 | 1.0 | 0.6 | 14877.0 | 47.0 | 108.3 | 67.4 | 8192.0 | 47.0 |
| | | 0.2 | 1.0 | 0.7 | 15264.3 | 11.8 | 25.7 | 13.4 | 7803.7 | 11.8 |
| | | 0.4 | 1.0 | 0.7 | 15633.0 | 14.0 | 45.0 | 34.1 | 7435.0 | 14.0 |
| | | 0.8 | 1.0 | 0.7 | 15459.3 | 42.2 | 17.7 | 4.9 | 7608.7 | 42.2 |

Table S16. Overview of the mapping statistics of generated samples in study case W4.

| | Sample index | SBS mutational signature | Chr1 copy number | Fraction mutated copies | % mutated positions | Raw reads Mean | Unmapped reads Mean |
|---|---|---|---|---|---|---|---|
| **Study case W4** | 1 | SBS2 | 2500 | 0.4 | 1.5 | 35M | 11684 |
| | 2 | SBS8 | | | | | 4144 |
| | 3 | SBS24 | | | | | 7459 |

Table S17. Overview of the percentage of covered and not covered exons in study case W4. The table shows the exon percentage covered with more than 10x, 50x, 100x, 200x and 300x as well as the percentage of uncovered exons grouped by the lengths 1-50 bp, and > 50 bp in samples in study case W4.

| | SBS mutational signature | Chr1 copy number | Fraction of mutated copies | Percentage of mutated positions | Percentage of exons covered with more then | | | | | Percentage of exons not covered | | |
|---|---|---|---|---|---|---|---|---|---|---|---|---|
| | | | | | | | | | | *Lengths* | | |
| | | | | | ***10x*** | ***50x*** | ***100x*** | ***200x*** | ***300x*** | ***All*** | ***1 - 50 bp*** | ***>50 bp*** |
| | | | | | Mean | Mean | Mean | Mean | Mean | Mean | Mean | Mean |
| **Study Case W4** | SBS2 | 2500 | 0.4 | 1.5 % | 100.0 | 100.0 | 100.0 | 99.6 | 92.4 | 0 | 0 | 0 |
| | SBS8 | | | | 100.0 | 100.0 | 100.0 | 99.6 | 92.6 | 0 | 0 | 0 |
| | SBS24 | | | | 100.0 | 100.0 | 100.0 | 99.6 | 92.5 | 0 | 0 | 0 |

Table S18. Overview of the inserted mutations in samples in study cases W4. The table exhibits the total number of unique mutations, mutation frequency in terms of the number of mutations per Mb DNA in an *in-silico* sample, and mean variant allele frequency (VAF).

| SBS mutational signture | Copy number | Fraction of mutated copies | Percentage of mutated positions | Number of unique mutations | Mutation frequency | VAF |
|---|---|---|---|---|---|---|
| SBS2 | 2500 | 0.4 | 1.5% | 52068 | 41.1 | 0.2 |
| SBS8 | | | | 52069 | 41.4 | 0.2 |
| SBS24 | | | | 52070 | 40.6 | 0.2 |
